# Low-latency neuromorphic closed-loop control of hippocampal ripples *in vivo*

**DOI:** 10.64898/2026.07.09.737518

**Authors:** Pedro Félix, Maria-Teresa Jurado-Parras, Joel Freitas, João Ventura, Liset M. de la Prida, Paulo Aguiar

## Abstract

Real-time closed-loop neuromodulation, in which stimulation is precisely timed to ongoing brain dynamics, holds transformative potential for treating neurological disorders and probing neural circuit function. However, it requires low-latency, energy-efficient processing of high-bandwidth neural signals that conventional computing architectures struggle to deliver. Neuromorphic computing, which emulates the event-driven and massively parallel operation of biological neural circuits, offers a compelling alternative. Yet, its integration into closed-loop frameworks validated in vivo for fast, transient oscillations has not been demonstrated. Here, we present a fully integrated neuromorphic framework for real-time detection and manipulation of hippocampal ripples: brief (30-100 ms), high-frequency (100-250 Hz) oscillations that are critical for memory consolidation and implicated in neurological disorders. We train compact spiking neural networks comprising 41 neurons and 530 parameters using surrogate-gradient backpropagation, achieving detection performance competitive with deep learning models across 23 recording sessions while consuming up to 200-fold less energy when deployed on SpiNNaker neuromorphic hardware. Integration with the open-source Open Ephys platform yields total closed-loop latencies of approximately 50 ms, enabling intra-event stimulation in up to 80% of ripples. Validating the complete sensing-processing-stimulation pipeline in awake, head-fixed mice, we demonstrate that neuromorphic-triggered optogenetic inhibition significantly alters ripple dynamics and reduces oscillatory energy. This work establishes a practical and accessible neuromorphic framework for low-latency closed-loop control of fast brain dynamics *in vivo*.

## 1. Introduction

Neuromodulation has become a powerful strategy for restoring or reshaping neural circuit activity in neurological disorders, including epilepsy and Parkinson’s disease (Davidson et al., 2024). Recent developments increasingly emphasize adaptive and personalized stimulation, driven by the integration of artificial intelligence into stimulation algorithms, the expansion of high-channel-count recording and stimulation interfaces, and the development of fully integrated neural devices (Oliveira et al., 2023; Sodagar et al., 2025; Yoo & Shoaran, 2021; B. Zhu et al., 2020). Within this context, closed-loop neuromodulation, in which stimulation is adjusted or triggered according to ongoing neural activity, has emerged as a central paradigm for both causal circuit interrogation and therapeutic intervention (Leite de Castro et al., 2024; Ramot & Martin, 2022; Sun & Morrell, 2014). By delivering stimulation only when relevant brain states or electrophysiological biomarkers are detected, closed-loop systems can improve specificity, reduce adverse effects, and lower power consumption relative to continuous open-loop stimulation (Soleimani et al., 2023; Sun & Morrell, 2014).

The effectiveness of closed-loop neuromodulation depends critically on timing. Neural biomarkers must be detected not only accurately, but also early enough for stimulation to interact with the underlying circuit dynamics. This imposes severe engineering constraints: high-bandwidth neural signals must be acquired, processed and converted into feedback commands with low latency, while operating within strict power and tissue-heating limits (Soleimani et al., 2023; Yoo & Shoaran, 2021). Although deep learning approaches have achieved impressive performance in neural signal decoding, their computational complexity and energy requirements often limit embedded deployment. Consequently, many implementations are forced to rely on external processors and wireless telemetry (Yoo & Shoaran, 2021; B. Zhu et al., 2020) - architectures that introduce latency overheads and energy demands often incompatible with intervention in short-lived neural events (Contreras, Truong, et al., 2024; Pawlak & Howard, 2025; B. Zhu et al., 2021). Therefore, there remains a pressing need for low-power, low-latency computational substrates capable of supporting real time, closed-loop operation at the edge.

Neuromorphic computing provides a promising solution to this challenge. Inspired by the event-driven, massively parallel and memory-efficient operation of biological neural circuits, neuromorphic systems are designed to process sparse temporal information with high energy efficiency (Indiveri & Liu, 2015; Kudithipudi et al., 2025; Schuman et al., 2017). These systems are commonly implemented using spiking neural networks (SNNs), which communicate through discrete spike events rather than continuous-valued activations (Eshraghian et al., 2023; Indiveri & Liu, 2015). Recent advances in surrogate-gradient learning and neuromorphic hardware have expanded the computational capabilities of SNNs, enabling compact networks to perform complex temporal pattern-detection tasks while retaining compatibility with specialized low-power processors (Eshraghian et al., 2023).

These properties have motivated growing interest in neuromorphic neural interfaces, leading to the development of promising systems relying on either custom-made or commercial neuromorphic hardware. Previous studies have explored neuromorphic approaches in epilepsy (seizure detection) (Bartels et al., 2025; Contreras, Huang, et al., 2024; Ronchini et al., 2021, 2023), as well as neuroprosthetics and brain-computer interfaces (BCIs) (Corradi & Indiveri, 2015; Dabbous et al., 2021; Dias et al., 2022; Shaikh et al., 2019). These advances reflect the mounting relevance and utility of neuromorphic systems, as the technology matures and increasingly approaches real-world and clinical applications (Contreras, Truong, et al., 2024; Covi et al., 2021; Donati & Indiveri, 2023; Pawlak & Howard, 2025; Yoo & Shoaran, 2021; B. Zhu et al., 2020; Y. Zhu et al., 2017). Of particular interest to this work are recent explorations of neuromorphic systems for detecting high-frequency oscillations (HFOs) in epilepsy (Sharifshazileh et al., 2020). **However, a complete neuromorphic closed-loop framework for real-time, state-dependent manipulation of fast transient brain activity patterns in vivo has not yet been demonstrated.** In particular, the field lacks accessible systems that combine live neuronal acquisition, neuromorphic event detection, and feedback stimulation using broadly adopted neuroscience infrastructure.

Hippocampal ripples provide a stringent test case for such systems. These are brief, high-frequency oscillations in the 100-250 Hz range, typically lasting approximately 30-100 ms, and forming part of the sharp-wave ripple complex (Buzsáki, 2015). Ripples play critical roles in memory consolidation and prospective replay and are increasingly implicated in pathological circuit states including epilepsy, Alzheimer’s disease and other neurological disorders (Aleman-Zapata et al., 2021; Lisgaras et al., 2025). While closed-loop manipulation of ripples has been essential for establishing causal links between these events and behavior or circuit function (Aleman-Zapata et al., 2021), their short duration and high frequency make them particularly demanding targets for real-time intervention: detection and stimulation must occur within tens of milliseconds, before the event has terminated. Ripples therefore provide both a biologically meaningful target and a rigorous benchmark for low-latency neuromorphic closed-loop control.

Here, we present a fully integrated neuromorphic framework for real-time detection and manipulation of hippocampal ripples in vivo. We trained compact SNNs on spike-encoded hippocampal local field potentials and benchmarked their performance against established conventional and deep learning ripple detectors. We then deployed the trained networks on SpiNNaker neuromorphic hardware (Painkras et al., 2013) and integrated them with the open-source Open Ephys platform (Siegle et al., 2017) to support real-time closed-loop operation. Through hardware-in-the-loop validation, we optimized the latency-performance trade-off and achieved closed-loop detection latencies of approximately 50 ms, enabling stimulation before ripple termination in up to 80% of events. Finally, using the complete sensing-processing-stimulation pipeline in awake, head-fixed mice, we demonstrate that neuromorphic-triggered optogenetic inhibition significantly alters ripple dynamics and reduces oscillatory energy. Together, these results validate the ability of neuromorphic systems to process live neural data, detect transient biomarkers, and trigger feedback stimulation within the strict timeframe required to manipulate fast brain dynamics in *vivo*.

## 2. Results

### 2.1. Spike-based encoding preserves ripple-band information for neuromorphic detection

To enable neuromorphic processing of hippocampal ripples, continuous local field potentials (LFPs) were first transformed into discrete spike trains. Raw hippocampal LFPs were bandpass filtered in the ripple range (100-250 Hz) and encoded into up (UP) and down (DN) spike events using an asynchronous delta-modulation scheme (Fig. 1A). This representation converts positive and negative signal deflections into separate event streams, providing a sparse spike-based input compatible with SNNs. The encoded spike trains were then downsampled from 30 kHz to 1 kHz to match the 1 ms simulation timestep used for neuromorphic deployment.

**Figure 1.**
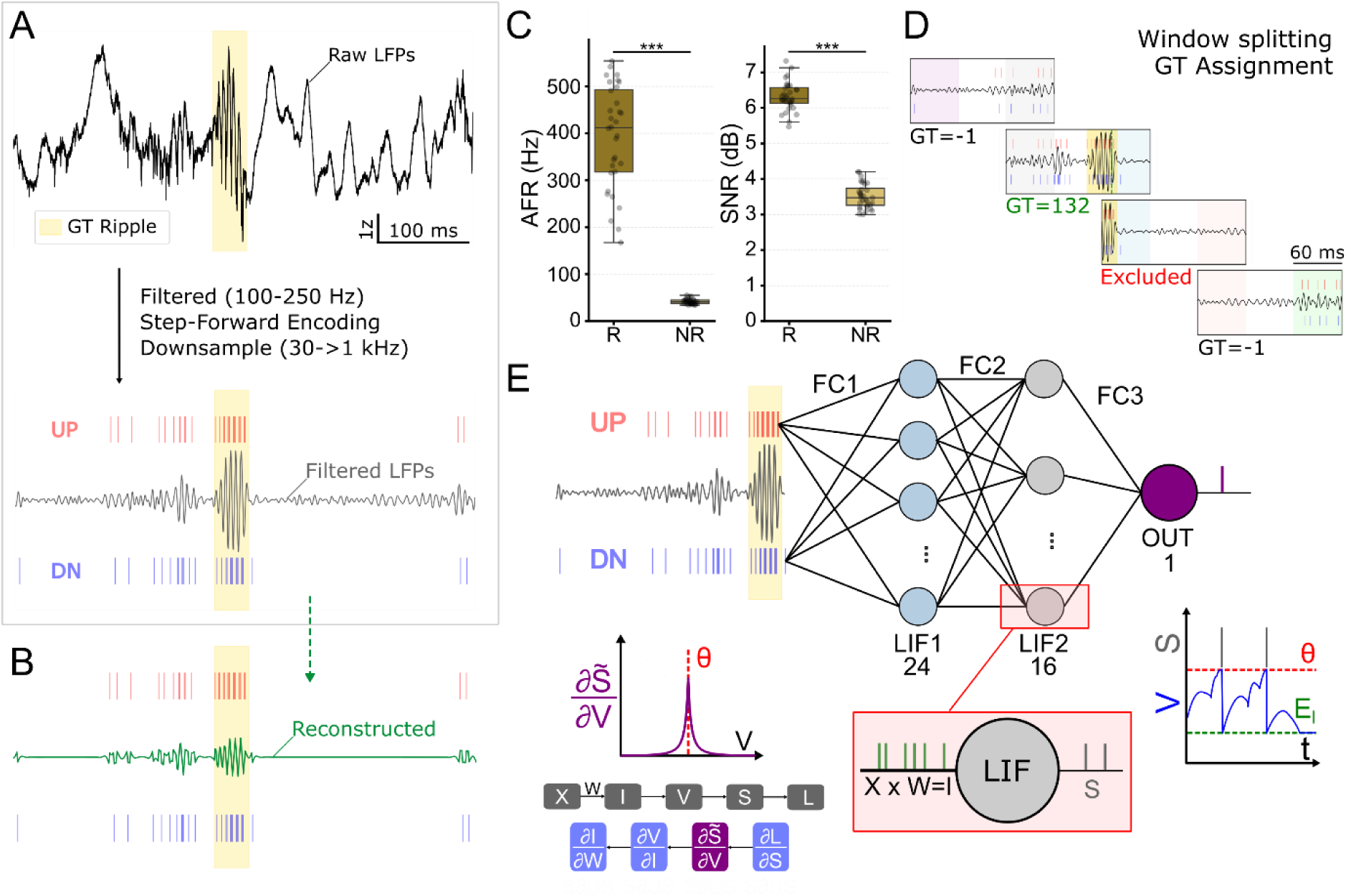
Spike-based encoding and SNN architecture for neuromorphic ripple detection. A, Hippocampal local field potentials (LFPs) were bandpass filtered in the ripple range (100-250 Hz) and converted into sparse UP and DN spike trains using asynchronous delta modulation. Encoded spike trains were downsampled from 30 kHz to 1 kHz to match the 1 ms timestep used for SNN simulation and neuromorphic deployment. B, Representative reconstruction of the ripple-band signal from UP/DN spike events, illustrating preservation of high-frequency ripple structure after spike-based encoding. C, Encoding metrics comparing ripple and non-ripple baseline segments. Average firing rate (AFR) was markedly higher during ripples than during baseline activity, and reconstructed signals showed higher signal-to-noise ratio (SNR) during ripple segments, indicating preferential encoding of event-relevant dynamics. Data are shown across channels/sessions as indicated in Methods; statistical comparisons were performed using two-sided Wilcoxon signed-rank tests. D, Continuous spike trains were segmented into overlapping windows for supervised training, with ground-truth labels assigned according to the temporal alignment between each window and annotated ripple events. E, SNN architecture used for ripple detection. The network receives two input streams corresponding to UP and DN events and comprises two fully connected hidden layers of leaky integrate-and-fire neurons, with 24 and 16 units, followed by a single output neuron that signals ripple detections. Networks were trained by backpropagation through time using surrogate-gradient learning.

We first assessed whether this encoding preserved the temporal structure required for ripple detection. Reconstruction of the filtered LFPs from the UP/DN spike trains reproduced the high-frequency oscillatory structure of ripple events, although with the expected attenuation associated with 1 kHz sampling and event-based quantization (Fig. 1B). Because the purpose of the encoding was event detection rather than waveform reconstruction, we quantified whether ripple periods were preferentially represented in the spike domain. UP/DN activity was markedly higher during ripples than during baseline non-ripple periods, with median average firing rates of 411.7 Hz during ripples compared with 40.8 Hz during baseline segments (Fig. 1C). Reconstruction quality was also higher during ripples, with median signal-to-noise ratios of 6.3 dB for ripple segments and 3.5 dB for baseline segments. Thus, the encoding selectively amplified event-relevant dynamics while maintaining a sparse representation of background activity.

The resulting spike trains were segmented into overlapping windows for supervised SNN training (Fig. 1D). We used a compact feed-forward architecture with two input channels corresponding to UP and DN events, two hidden layers of leaky integrate-and-fire (LIF) neurons comprising 24 and 16 units, and a single output neuron acting as the ripple detector (Fig. 1E). Networks were trained by backpropagation through time using surrogate gradients (SG-BPTT) to overcome the non-differentiability of spike generation. This architecture contains only 41 neurons and 530 trainable parameters, providing a deliberately minimal model for testing whether ripple-relevant temporal structure could be detected by a hardware-compatible SNN.

### 2.2. Compact SNNs detect ripples with performance comparable to larger machine-learning detectors

We next evaluated whether the trained SNNs could detect hippocampal ripples in continuous LFP recordings. Performance was assessed across 23 recording sessions from five mice, comprising 2719 manually annotated ripple events. Representative continuous recordings showed that output spikes were temporally aligned with annotated ripples, while also revealing occasional false positives and missed low-amplitude events (Fig. 2A). Inspection of full-network spiking activity during continuous playback further showed that detections emerged from sparse and structured activity across the hidden layers (Fig. 2B). Several hidden units remained largely silent, whereas a small subset of neurons consistently preceded output spikes, suggesting that the network relies on a compact and partially redundant representation that may be further reducible by pruning.

**Figure 2.**
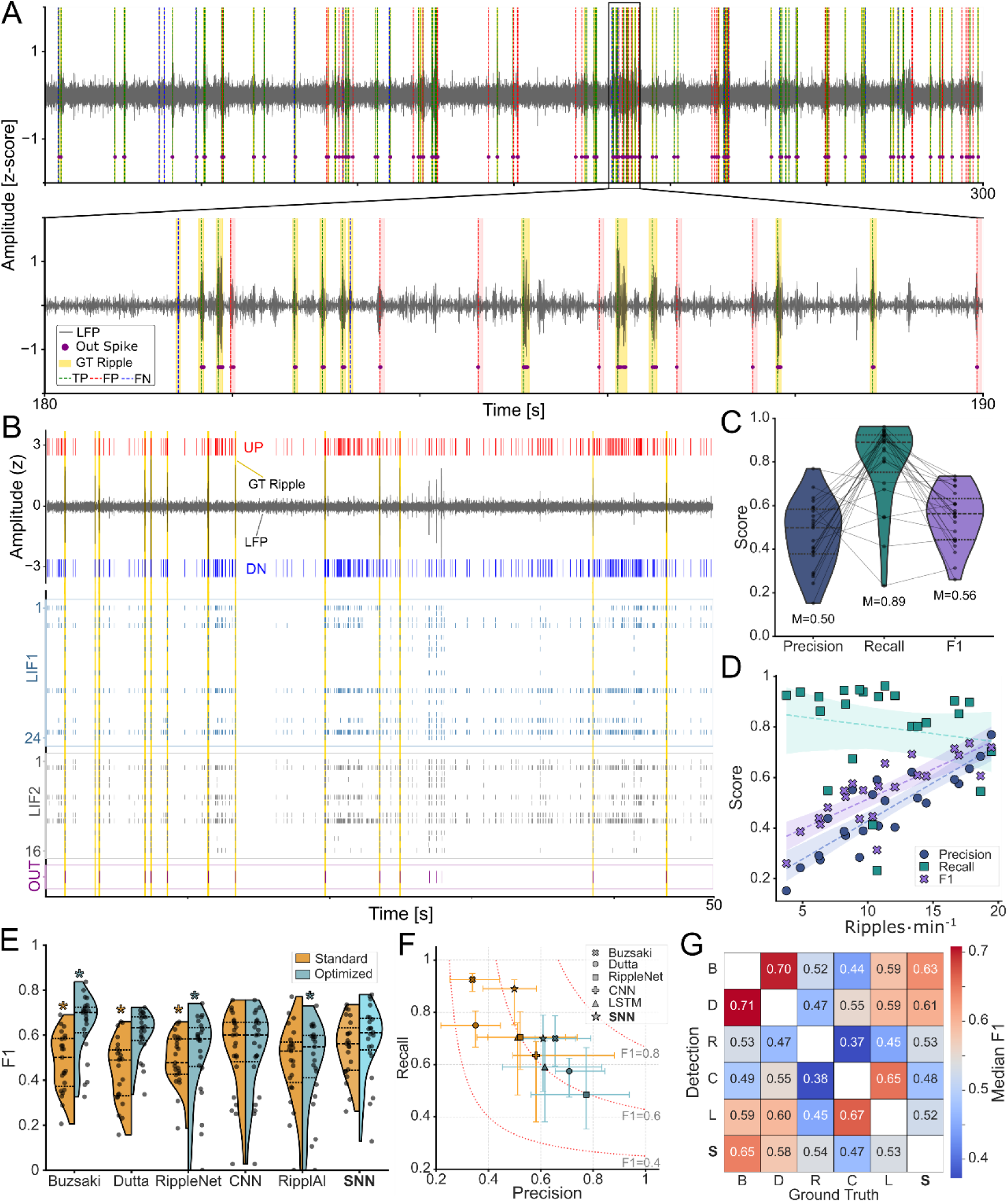
SNN ripple detection performance in continuous LFPs and comparison with established detectors. A, Representative SNN detection over 300 s of continuous hippocampal LFPs, with a 10 s zoomed segment. Ripple-band LFPs, manually annotated ground-truth (GT) ripples, SNN output spikes and event classifications are shown. TP, true positive; FP, false positive; FN, false negative. B, Representative raster plot showing UP/DN input events, GT ripples, ripple-band LFPs, hidden-layer spiking activity and output spikes during continuous SNN processing. Output spikes correspond to putative ripple detections. C, Distribution of precision, recall and F1-score across recording sessions (n = 23), aggregated as the median performance of the selected SNN configurations. The SNNs showed high recall and lower precision, reflecting the precision-recall trade-off observed across sessions. D, Relationship between ripple rate and detection performance. Ripple rate was strongly correlated with precision and F1-score (Spearman’s ρ = 0.85, p < 0.01 for both), whereas recall remained stable across sessions (ρ = −0.15, p = 0.48). E, Benchmark comparison against conventional filter-based detectors and deep-learning models: Buzsáki-style RMS detector (Fernández-Ruiz et al., 2019), Dutta filter-based detector (Dutta et al., 2018), RippleNet (Hagen et al., 2021), 1D-CNN (Navas-Olive et al., 2022), and LSTM model (Navas-Olive et al., 2024). Models were evaluated using standard out-of-the-box parameters and parameters optimized for maximum F1-score on this dataset. The SNN achieved performance comparable to larger deep-learning models, while the optimized Buzsáki-style detector reached the highest absolute F1-score. F, Precision-recall representation of SNN and benchmark detector performance under standard and optimized configurations. G, Inter-detector consistency matrix. Similarity was quantified as the median F1-score obtained when using each optimized detector’s predictions as pseudo-ground truth for the others. The SNN showed strongest agreement with the optimized Buzsáki-style detector and broader convergence with filter-based approaches.

Across recording sessions, the SNNs detected ripples with high sensitivity but lower selectivity, consistent with the known difficulty of ripple detection in noisy biological recordings. Median recall was 0.89 (IQR, 0.75-0.92), whereas median precision was 0.50 (IQR, 0.38-0.58), yielding a median F1-score of 0.56 (IQR, 0.44-0.63; Fig. 2C and Table S2). This precision-limited profile is consistent with previous ripple and high-frequency oscillation detectors, where visually annotated events are difficult to distinguish from sharp transients, muscle artifacts and irregular high-frequency activity with overlapping spectral content (Hagen et al., 2021; Liu et al., 2022; Roux et al., 2017; Zelmann et al., 2012; Zhou et al., 2022). Substantial inter-session variability was also observed, reflecting the characteristic heterogeneity of biological recordings and the precision-recall trade-off commonly reported for ripple detectors (Navas-Olive et al., 2024).

To better understand this variability, we examined the relationship between ripple rate and detection performance (Fig. 2D). Ripple rate was strongly associated with both precision and F1-score (Spearman’s ρ = 0.85, p < 0.01 for both), whereas recall remained comparatively stable across sessions (ρ = −0.15, p = 0.48). Thus, the SNNs retained sensitivity across heterogeneous physiological states, but precision decreased in recordings with fewer ripples, where false positives have a proportionally larger impact on aggregate F1-score.

We then benchmarked the SNNs against established conventional and deep learning-based ripple detectors, including Buzsáki-style RMS detector (Fernández-Ruiz et al., 2019), the Dutta filter-based detector (Dutta et al., 2018), RippleNet (Hagen et al., 2021), a 1D-CNN (Navas-Olive et al., 2022), and an LSTM model, RipplAI (Navas-Olive et al., 2024) (Fig. 2E-G, Table S4 and Fig. S3). The SNNs achieved performance comparable to larger deep learning architectures. When the best-performing SNN was compared with optimized models, it reached a median F1-score of 0.61 (IQR, 0.53-0.67), significantly outperforming RippleNet (p = 0.012) and the LSTM model (p = 0.013), while performing comparably to the 1D-CNN (p = 0.22).

The highest absolute F1-score was obtained by the optimized Buzsáki-style RMS detector (median F1 = 0.70; IQR, 0.61-0.72), which outperformed the SNN and the other machine-learning models (p < 0.01). This confirms that classical filter-based approaches remain highly effective when carefully calibrated to a specific dataset. However, the comparison between standard and optimized detector configurations revealed an important operational distinction. When used with standard, out-of-the-box parameters, conventional filter-based detectors showed marked performance degradation, whereas the SNNs maintained competitive performance and significantly outperformed both standard Buzsáki and Dutta detectors (p < 0.01). This supports the suitability of the SNN approach for real-time closed-loop settings, where extensive post-hoc parameter tuning is impractical.

Precision-recall analysis further illustrated this trade-off (Fig. 2F). Standard detector configurations generally favoured sensitivity, whereas F1-optimized configurations improved precision at the cost of recall. In this space, both the optimized Buzsáki detector and the SNN occupied comparatively balanced operating regimes, closer to the upper-right region of the precision-recall plot than most other approaches. Finally, inter-detector consistency analysis showed that SNN predictions were most closely aligned with the optimized Buzsáki detector (relative F1 = 0.65) and more broadly with filter-based methods (Fig. 2G). This suggests that the SNN captures core ripple-band features similar to those used by conventional detectors, while retaining a compact architecture suitable for neuromorphic hardware deployment.

### 2.3. Detection errors arise from overlapping low-amplitude ripple-band events

To characterize the operational limits of the SNN ripple detectors, we compared true positives (TPs), false positives (FPs) and false negatives (FNs) using ripple-band signal features (Fig. 3). Visual inspection of representative 100-250 Hz LFP snippets showed that TPs typically corresponded to prominent ripple oscillations, whereas FPs and FNs often appeared as lower-amplitude ripple-like events (Fig. 3A). This pattern was confirmed by event-triggered ripple-band power profiles: TPs showed substantially higher power around the event peak, while FP and FN profiles were markedly similar and lower in amplitude (Fig. 3B).

**Figure 3.**
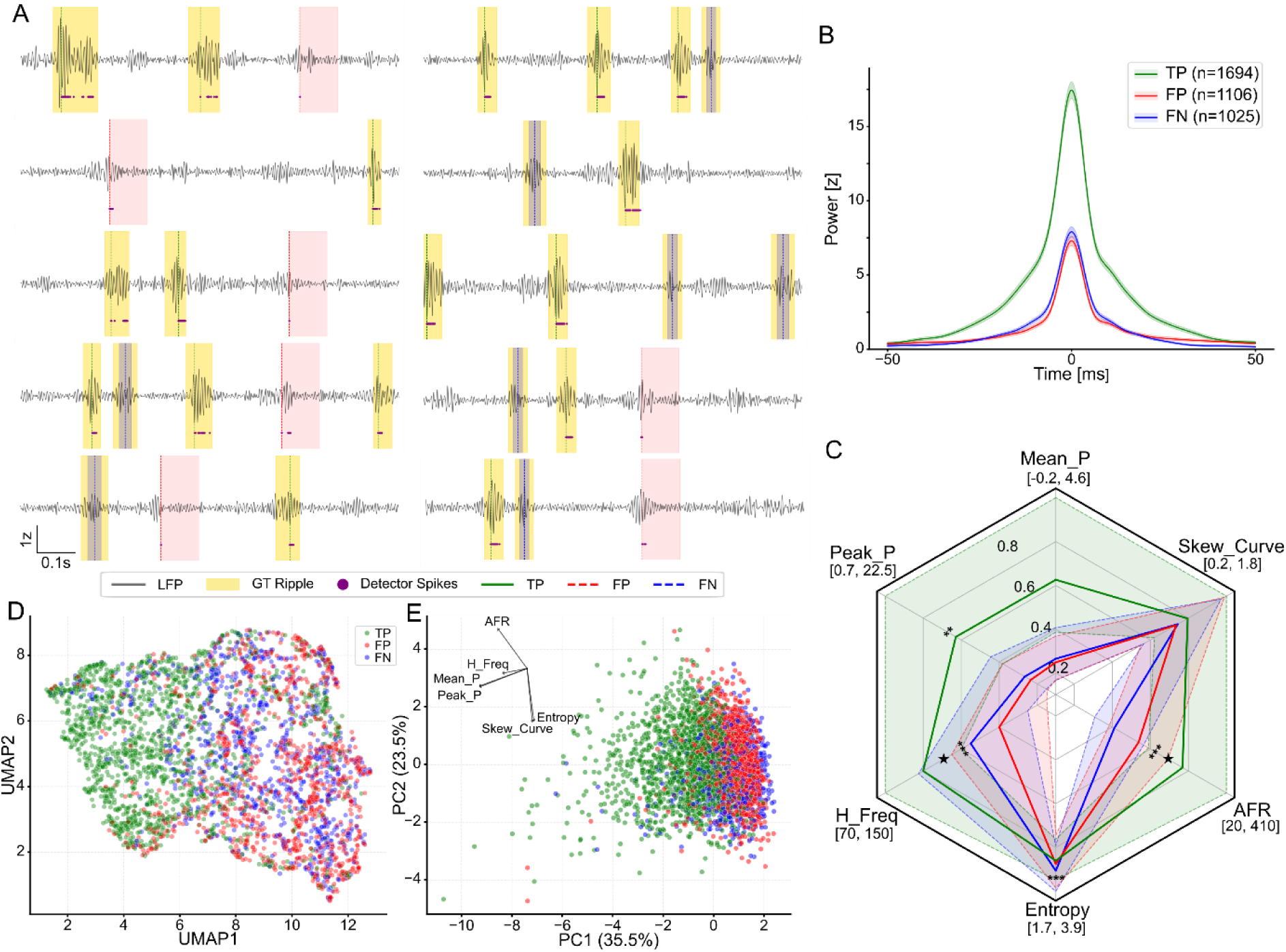
Feature structure of SNN-detected, missed and false-positive ripple-band events. A, Representative 400 ms LFP snippets filtered in the ripple band (100-250 Hz), illustrating true positives (TPs), false positives (FPs) and false negatives (FNs). Manually annotated ground-truth ripples and SNN output spikes are indicated. B, Event-triggered ripple-band power centered on the event peak (±50 ms). TPs showed higher ripple-band power, whereas FPs and FNs exhibited similar lower-amplitude profiles. Data are shown as mean ± 95% confidence interval for events detected or missed by the best-performing network. C, Feature profiles of TP, FP and FN events. Radar plots show median and interquartile range across events for peak power, mean power, average UP/DN firing rate (AFR), spectral entropy, peak frequency and skewness of the curvature. Features were normalized for visualization. TPs showed higher power, AFR, peak frequency and curvature skewness than both FPs and FNs. Direct comparison between FPs and FNs showed that FPs had lower peak frequency (Mann-Whitney test, p < 0.001, Cliff’s δ = 0.2) and higher AFR (p < 0.001, Cliff’s δ = −0.3). Differences in entropy and peak power reached statistical significance but had negligible effect sizes (Cliff’s δ ≤ 0.1). D, Uniform Manifold Approximation and Projection (UMAP) visualization of the event-feature space. E, Principal Component Analysis (PCA) visualization of the same feature space. Both projections revealed a continuous distribution without clear separation between event classes, with substantial overlap between FP and FN events.

We next quantified electrophysiological features associated with each event class, including ripple-band power, peak frequency, spectral entropy, spike-encoding activity and waveform curvature (Fig. 3C and Table S5). As expected, TPs showed higher power, peak frequency and UP/DN firing rate than both FPs and FNs (p < 0.001, Cliff’s δ > 0.35), consistent with their stronger oscillatory structure and more reliable spike-domain representation. Direct comparison between FPs and FNs revealed only modest differences. FPs had a lower peak frequency than FNs (p < 0.001, Cliff’s δ = 0.2), suggesting that some false detections may reflect lower-frequency events or artifacts entering the ripple band. FPs also showed higher UP/DN average firing rate (p < 0.001, Cliff’s δ = −0.3), indicating that denser bursts of encoded input spikes can occasionally drive the output neuron despite the absence of a ground-truth annotation.

Despite these statistically significant differences, no individual feature reliably separated the three event classes. Dimensionality-reduction analyses using UMAP and PCA revealed a continuous feature space without clearly separable clusters (Fig. 3D,E). The strongest overlap occurred between FPs and FNs, indicating that many errors arise from ambiguous low-amplitude events near the boundary of manual ripple annotation. This overlap also argues against the SNN operating as a simple power threshold or spike counter. Instead, the network appears to integrate amplitude, frequency and temporal spike-pattern information, but remains constrained by the intrinsic ambiguity of ripple-band events in continuous biological recordings.

### 2.4. Neuromorphic deployment preserves ripple detection while reducing energy consumption

After validating the SNNs in software, we deployed a selected network on SpiNNaker neuromorphic hardware and integrated it with Open Ephys to evaluate real-time closed-loop operation (Fig. 4A). In this configuration, pre-recorded LFPs were streamed through the Open Ephys graphical user interface (GUI), encoded into UP/DN spikes and processed by the SNN running on the SpiNNaker board. Feedback events generated by the output neuron were returned to Open Ephys and logged for post-hoc comparison with manually annotated ripples. This setup allowed us to quantify how neuromorphic deployment affected detection performance before proceeding to full hardware-in-the-loop latency validation.

**Figure 4.**
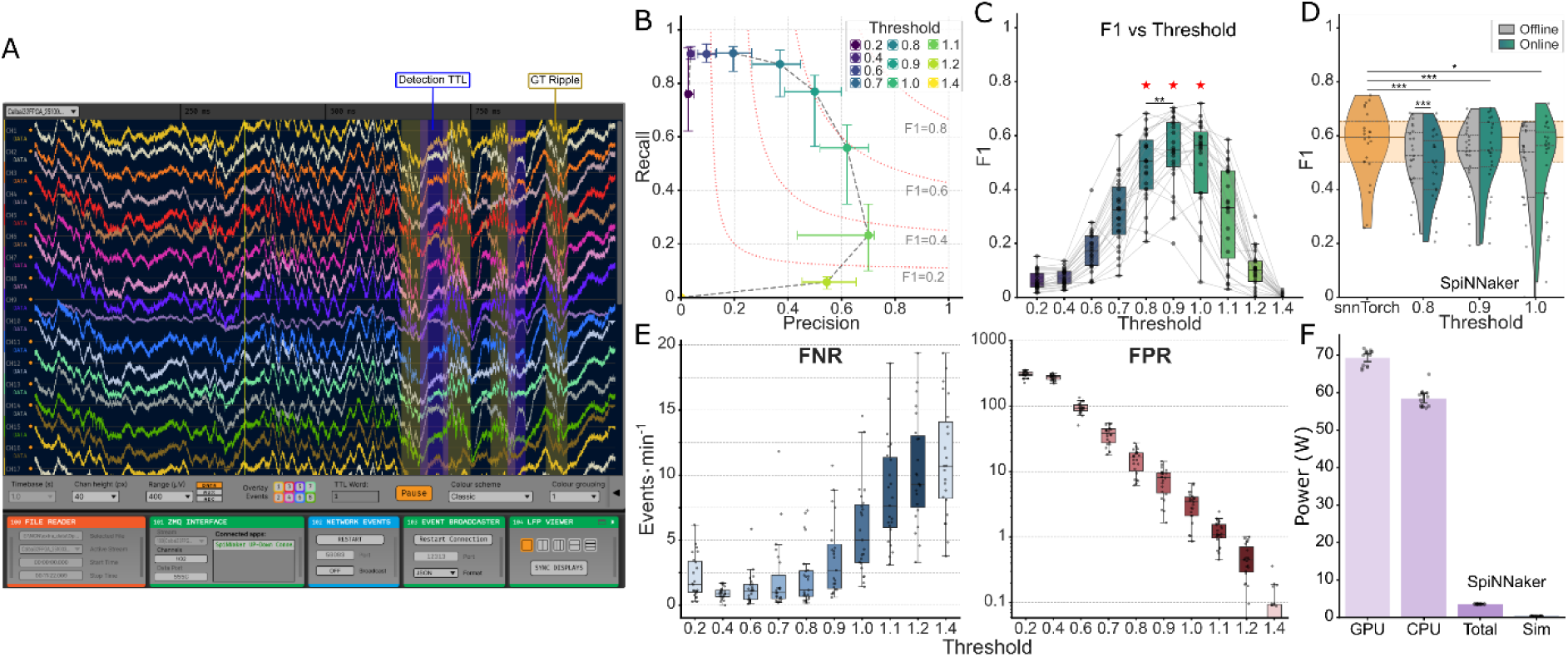
Neuromorphic deployment preserves online ripple detection and enables energy-efficient operation. A, Open Ephys signal chain used for online in silico validation. Pre-recorded LFPs were streamed through the Open Ephys File Reader and ZMQ Interface plugins to an external Python bridge, encoded into UP/DN spikes and processed by the SNN running on SpiNNaker hardware. Feedback events generated by the output neuron were returned to Open Ephys through Network Events and logged for post-hoc analysis. B, Precision-recall trade-off across output-neuron firing thresholds during SpiNNaker operation using 6 ms buffers. Data points show median performance across recording sessions (n = 23), with error bars indicating interquartile range. C, F1-score as a function of firing threshold. Thresholds between 0.8 and 1.0 produced the highest overall performance and significantly outperformed the remaining tested thresholds after Holm-Bonferroni correction (Wilcoxon signed-rank tests, p < 0.01). Within this range, threshold 0.9 achieved higher F1-score than threshold 0.8 (p < 0.01). D, Performance impact of deployment from software simulation to SpiNNaker hardware and transition from offline continuous processing to online buffered processing. The software-to-hardware transition produced a modest but significant reduction in F1-score at thresholds 0.8 (p < 0.001), 0.9 (p < 0.001) and 1.0 (p < 0.05). In contrast, online buffered processing preserved performance relative to offline SpiNNaker processing at the most balanced operating thresholds. E, False positive and false negative rates as a function of firing threshold. Increasing the threshold reduced false positives per minute while increasing missed detections, illustrating the operational trade-off between stimulation specificity and detection sensitivity. F, Power consumption (Watts) associated with SNN simulation and inference using a regular GPU, CPU, and the SpiNNaker neuromorphic board (total power and power dedicated to simulation), n=14 sessions. SpiNNaker shows a 16/20-fold reduction in power consumption compared to CPU/GPU, that can go up to 170/200-fold when considering only the power consumed by the actual simulation (disregarding background processes).

We first evaluated the effect of firing threshold on online SpiNNaker performance using 6 ms LFP buffers (Fig. 4B,C and Table S6). Varying the threshold produced the expected precision-recall trade-off. Lower thresholds increased sensitivity but generated more false positives, whereas higher thresholds improved selectivity at the cost of missed events. Thresholds between 0.8 and 1.0 provided the best overall F1-scores, all significantly outperforming the remaining tested thresholds after Holm-Bonferroni correction (p < 0.01). Within this range, threshold 0.8 maximized recall (median = 0.87) but yielded lower precision (median = 0.37), whereas threshold 1.0 increased precision (median = 0.62) but reduced recall to 0.56. A threshold of 0.9 provided the most balanced operating point, combining robust recall (median = 0.77) with moderate precision (median = 0.50).

Because false detections translate directly into unnecessary stimulation in a closed-loop system, we also quantified error rates in events per minute (Fig. 4E). Increasing the firing threshold reduced the false positive rate (FPR) while increasing the false negative rate (FNR). At threshold 0.8, the SNN generated a median of 16.9 FPs per minute and 1.2 FNs per minute. Threshold 0.9 reduced FPs to 8.1 per minute while increasing FNs to 2.7 per minute. Threshold 1.0 further reduced FPs to 3.5 per minute but increased FNs to 5.0 per minute. Thus, threshold selection provides a direct way to tune the detector according to the target application: sensitivity can be prioritized when sensitivity or early intervention is critical, whereas specificity can be prioritized when minimizing unnecessary stimulation is more important.

We then quantified the performance impact of two operational transitions: deployment from software simulation to SpiNNaker hardware, and transition from offline continuous processing to online buffered processing (Fig. 4D and Table S7). The software-to-hardware transition produced a modest reduction in F1-score, with median decreases of 15.6%, 8.0% and 5.2% at thresholds 0.8, 0.9 and 1.0, respectively. These losses likely reflect differences between the discrete-time snnTorch training environment and the continuous-time PyNN implementation running on SpiNNaker (Pedersen et al., 2024; Schuman et al., 2017). In contrast, the transition from offline to online operation within the SpiNNaker environment had minimal additional impact. Offline and online performance were statistically indistinguishable at threshold 0.9 (−0.5% difference, p = 0.08) and threshold 1.0 (−4.3% difference, p = 0.15), with only threshold 0.8 showing higher offline performance (4.7%, p < 0.01). Moreover, a linear mixed-effects model accounting for inter-session variability indicated that online SpiNNaker performance at threshold 0.9 remained statistically comparable to the original software baseline (coefficient = −0.30, p = 0.15). These results show that although neuromorphic deployment introduces a measurable software-to-hardware gap, the integrated online processing pipeline preserves useful ripple-detection performance.

Finally, we quantified energy consumption by simulating SNN inference on CPU, GPU and SpiNNaker over a subset of the sessions (n=14, Fig. 4F). GPU-and CPU-based simulation consumed medians of 70 and 53 W and failed to operate in real time. In contrast, the SNN deployed on SpiNNaker operated in biological real time, processing the sessions without delays while consuming ≈3.6 W at board level, of which approximately 0.35 W was attributed to the simulation. This corresponds to an estimated 16/20-to 170/200-fold reduction in power consumption relative to GPU/CPU-based simulation, depending on whether total board power or the simulation-attributed power is considered. Together, these results demonstrate that compact SNNs can be transferred to neuromorphic hardware, preserve online ripple-detection performance and operate in real time with substantially reduced energy consumption.

### 2.5. Full closed-loop system allows low latencies compatible with ripple intervention

After confirming that SNN deployment on SpiNNaker preserved online detection performance, we quantified the latency of the complete closed-loop pipeline under realistic hardware conditions. Pre-recorded hippocampal LFPs were streamed through a National Instruments data acquisition board to emulate live acquisition by the Open Ephys system (Fig. 5A). This hardware-in-the-loop configuration allowed us to measure round-trip latency across the full acquisition-processing-feedback pathway, including data acquisition and buffering, spike encoding, SpiNNaker processing and feedback triggering.

**Figure 5.**
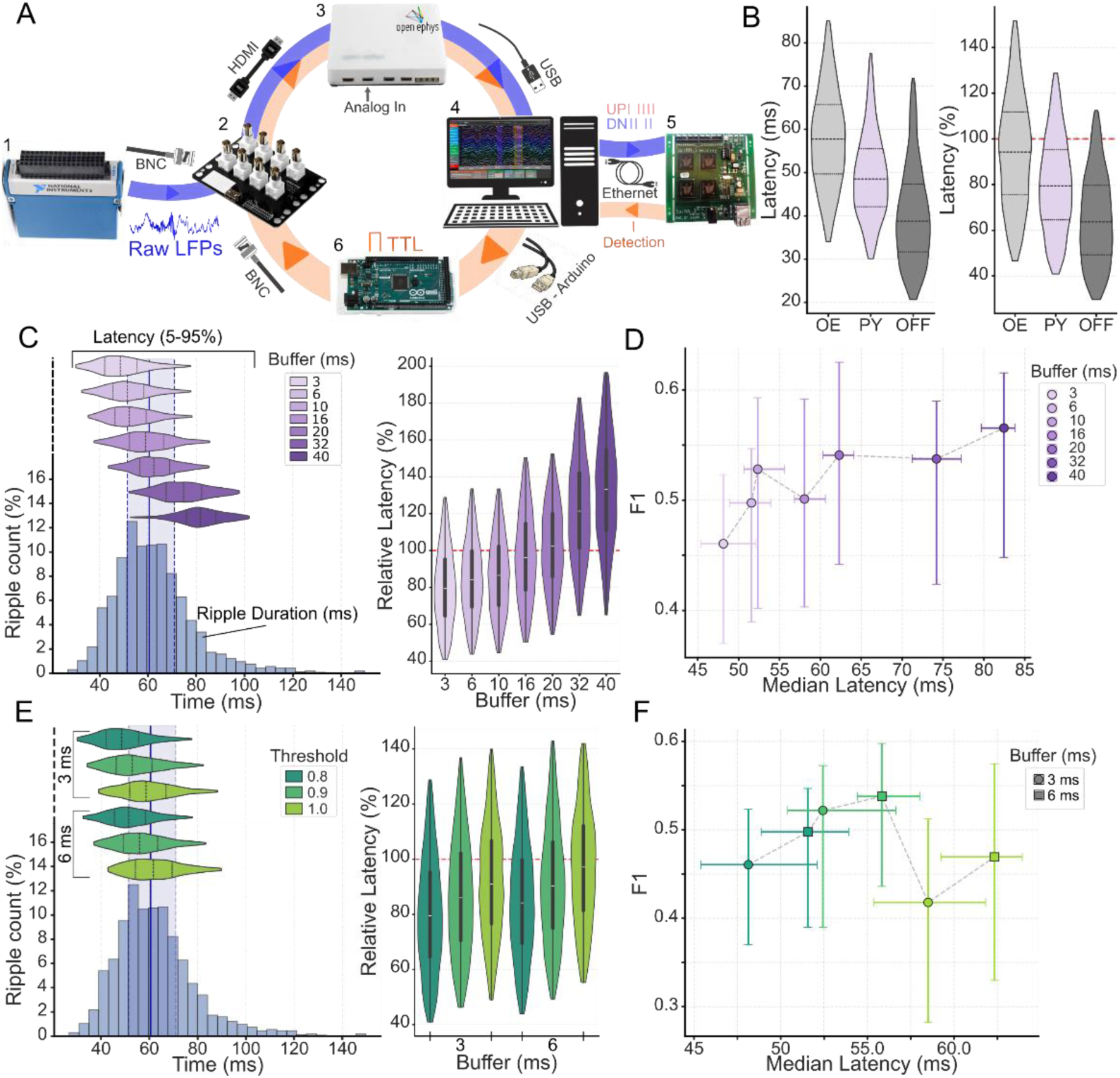
Hardware-in-the-loop validation of closed-loop latency and intervention timing. A, Hardware validation setup. Pre-recorded hippocampal LFPs were generated by a National Instruments NI-9264 data acquisition board (1), routed through the Open Ephys I/O board (2) and acquired by the Open Ephys Acquisition Board (3). Signals were streamed to the host computer (4), encoded into UP/DN spikes and sent through a Python bridge to the SNN running on SpiNNaker hardware (5). SNN detections triggered an Arduino MEGA 2560 output (6), which was looped back to the Open Ephys I/O board for round-trip latency measurement. B, Effect of feedback-triggering pathway on round-trip latency. Latency was compared using the Open Ephys Arduino Output plugin, direct Python control of the Arduino output and the offline simulation baseline, with buffer size fixed at 3 ms and firing threshold fixed at 0.8. Direct Python control significantly reduced latency relative to the Open Ephys plugin (Mann-Whitney U, p < 0.001). Data is shown as median with 5th-95th percentiles. C, Relationship between buffer size, round-trip latency and ripple duration. Latency increased approximately linearly with buffer size, as quantified by a linear mixed-effects model (slope = 0.84 ms per 1 ms buffer increase, p < 0.001). Relative latency analysis showed that stimulation before ripple termination was consistently achieved only with short buffers. D, Latency-performance trade-off across buffer sizes. Median F1-score and latency are shown as a function of buffer size. Larger buffers modestly improved detection performance but imposed a substantial latency cost. F1-score was strongly correlated with buffer size (Spearman’s ρ = 0.93, p < 0.01; linear mixed-effects model predictor = 0.002, p < 0.001). Data are shown as median ± interquartile range. E, Effect of firing threshold on latency using the two shortest buffer configurations, 3 ms and 6 ms. Higher thresholds increased latency, although all tested configurations maintained median relative latencies below ripple duration. F, Joint latency-performance trade-off across buffer size and firing threshold. Increasing the buffer from 3 to 6 ms improved F1-score across thresholds (Wilcoxon signed-rank test, p < 0.01). Threshold 0.9 yielded the highest median F1-score and significantly outperformed threshold 1.0 (Wilcoxon signed-rank test, p < 0.001), but was not statistically superior to threshold 0.8, which provided lower latency and higher intra-event intervention probability.

We first evaluated the feedback-triggering method using a 3 ms buffer and a firing threshold of 0.8 (Fig. 5B). Direct control of the Arduino output through the Python bridge significantly reduced round-trip latency relative to the Open Ephys Arduino Output plugin, with a median reduction of 9.2 ms (95% CI, 8.2-10.1 ms; p < 0.01, Cliff’s δ = −0.4). Comparing this optimized hardware configuration with the in-silico baseline indicated a hardware communication delay of approximately 9.8 ms (median; 95% CI, 8.9-10.6 ms), consistent with previously reported USB communication delays in Open Ephys-based closed-loop systems (Dutta et al., 2018). This reduction had a direct functional impact: direct Python triggering enabled stimulation before ripple termination in approximately 80% of events, whereas routing feedback through the Open Ephys plugin reduced this proportion to 59%. Thus, millisecond-scale optimization of the feedback path substantially improved the probability of intra-event intervention.

We next quantified the effect of Open Ephys buffer size on latency and detection performance using a fixed firing threshold of 0.8 (Fig. 5C,D and Table S8). Round-trip latency increased nearly linearly with buffer size. A linear mixed-effects model accounting for inter-session variability estimated the relationship as Latency (ms) = 47.45 + 0.84 × Buffer, indicating that each 1 ms increase in buffer duration added approximately 0.84 ms to the median latency. The minimum latency was obtained with a 3 ms buffer (median = 48.53 ms; IQR, 41.37-56.52 ms). To achieve stimulation before ripple termination in more than 75% of events, buffer sizes had to remain below 10 ms.

Increasing buffer size modestly improved detection performance, primarily through a higher precision. Median F1-score increased from 0.47 with a 3 ms buffer up to 0.57 with a 40 ms buffer (Spearman’s ρ = 0.93, p < 0.01; LMM predictor = 0.002, p < 0.001). This likely reflects reduced temporal fragmentation, as longer buffers provide more temporal context for distinguishing ripple oscillations from background activity. However, the effect of buffer size on performance was small compared with inter-session variability. Buffer size accounted for only 2.7% of F1-score variance (marginal R² = 0.027), whereas inter-session variability accounted for more than 90% of the variance (conditional R² = 0.943). Thus, larger buffers provide only limited performance gains while imposing a substantial latency cost.

Having established that short buffers are required for real-time ripple intervention, we evaluated the interaction between buffer size and firing threshold using the two fastest buffer configurations, 3 ms and 6 ms, and the three best-performing thresholds identified prior, 0.8, 0.9 and 1.0 (Fig. 5E,F and Table S9). Both buffer duration and firing threshold independently increased latency. A linear mixed-effects model confirmed positive effects of threshold (coefficient = 52.59, z = 27.6) and buffer size (coefficient = 0.87, z = 8.8) on round-trip latency. The lowest latencies were obtained with threshold 0.8, but reducing the buffer could compensate for the latency cost of a higher threshold: the latency difference between a 6 ms buffer at threshold 0.8 and a 3 ms buffer at threshold 0.9 was only 1.47 ms (95% CI, 0.63-2.47 ms; Cliff’s δ = 0.096).

Threshold 1.0 was not suitable for real-time ripple intervention because its higher selectivity came at the cost of reduced sensitivity and increased latency. With a 3 ms buffer, threshold 1.0 yielded a median latency of 58.63 ms (IQR, 51.05-67.67 ms), missing approximately half of all true ripples and achieving intra-event stimulation in less than 65% of events. In contrast, thresholds 0.8 and 0.9 provided viable operating regimes. Threshold 0.8 minimized latency and maximized recall, enabling intra-event stimulation in 80.4% of ripples with a 3 ms buffer and 75.4% with a 6 ms buffer. Threshold 0.9 improved overall F1-score by increasing precision, achieving median F1-scores of 0.52 and 0.54 with 3 ms and 6 ms buffers, respectively, while still allowing stimulation before ripple termination in 72.1% and 66.3% of events.

Together, these results define a practical latency-performance operating space for neuromorphic closed-loop ripple intervention. The fastest configuration, using a 3 ms buffer and threshold 0.8, achieved median closed-loop latencies of approximately 50 ms and maximized the probability of stimulation within the ripple duration. A more selective configuration, using threshold 0.9 and a 6 ms buffer, improved detection performance at the cost of an approximately 7 ms latency increase. Thus, the neuromorphic pipeline can be tuned either for maximal intervention speed and sensitivity or for improved stimulation specificity, depending on the experimental objective.

### 2.6. Neuromorphic System achieves real-time ripple manipulation *in vivo*

Following thorough validation of our system in simulated environments (in silico, hardware-in-the-loop), we proceeded to validate the framework in a real-world, *in vivo* ripple manipulation task. We tested the system across twelve sessions (4 OFF, 8 ON) from two mice (total of 1699 post-hoc ripple events). The closed-loop pipeline consisted of recording raw LFPs from the CA1 hippocampus of awake, head-fixed mice using high-density µLED optoelectrodes. Signals were acquired via the Open Ephys board and pre-processed and routed via a Python bridge running on the host computer to the SNNs running on SpiNNaker. Upon detection, the system triggered an Arduino MEGA 2560 to drive optogenetic stimulation of inhibitory neurons (Figure 6A).

**Figure 6.**
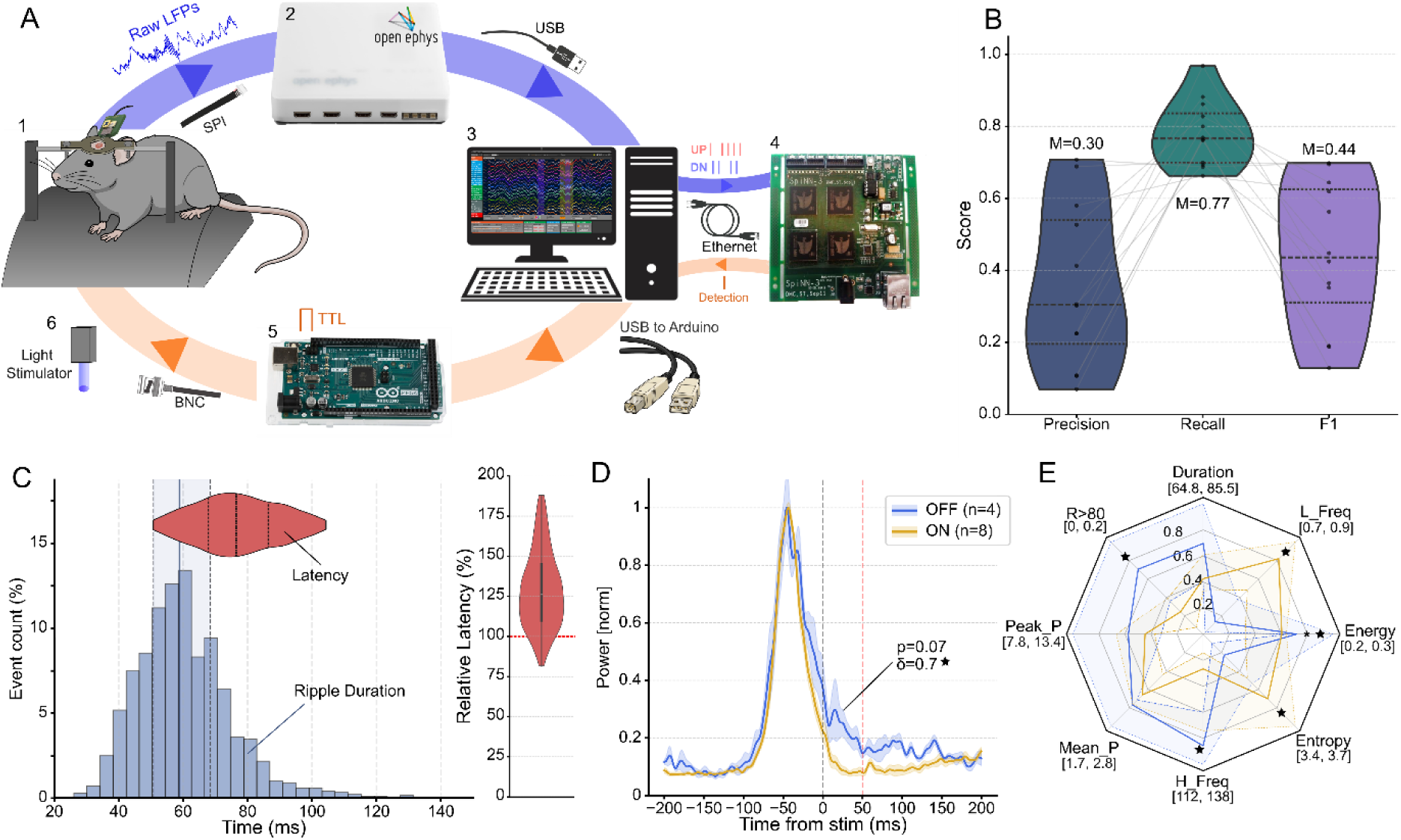
In vivo closed-loop ripple manipulation using the neuromorphic framework. A. Schematic of the closed-loop optogenetics setup. Neural signals from the CA1 hippocampus of a head-fixed mouse (1) are acquired via µLED optoelectrodes connected to an Open Ephys Acquisition Board (2). Data is streamed to the host computer (3), and spike-encoded for processing by the SNNs running on the SpiNNaker board (4). Detections trigger an Arduino MEGA 2560 (5) to drive a 100 ms inhibitory optogenetic pulse via the light stimulator (6). B. In vivo detection performance. Post-hoc evaluation of performance across sessions (n=12) displays wide inter-session variability. C. Round-trip latency versus event duration. Utilizing a preliminary unoptimized configuration (20 ms buffer), the median detection latency exceeded the median event duration (median relative latency = 126%). D. Peri-detection power profile. Average ripple-band power (mean ± SEM) centered on detection time (t=0), normalized per window. Comparison of mean power in the post-detection window (+50 ms) reveals a nonsignificant but large magnitude reduction in “ON” sessions (4 sessions OFF, 8 sessions ON, Mann-Whitney U, p=0.07, δ=0.69). E. Radar Plot characterizing the impact of light stimulation on ripple features (top 25% longest events per session). Data is shown as the median and interquartile range across sessions. Features include: Peak and Mean Power (z-scored), Duration (ms), Spectral Entropy, Peak Frequency (H_Freq, Hz), ratio of long ripples (R>80 ms, calculated across all ripples), low frequency contribution (L_Freq) and Energy. Values are normalized between the minimum and the 3rd quartile for visualization. Optogenetic inhibition seems to have significantly altered ripple dynamics, as “ON” sessions exhibit significantly lower energy (*p* = 0.048*, δ=0.75), and also higher spectral entropy (δ=-0.56), lower peak frequency (δ=0.56), a reduced fraction of long-duration ripples (δ=0.56), and a higher low frequency contribution (δ=-0.69).

Configured to optimize sensitivity, the SNN detectors achieved a median F1-score of 0.44 and successfully detected a median of 77% of all ground-truth ripples (Figure 6B). It should be noted that these experiments were conducted using a preliminary configuration, prior to latency optimization (i.e. with a buffer of 20 ms and OE plugin output), which led to a high median latency of 76.47 ms (IQR 66.67-88.13), leading to only 13.7% of ripples receiving stimulation before termination, compromising real-time intervention (Figure 6C).

Despite this limitation, evaluating peri-detection ripple power revealed an effect of optogenetic stimulation (Figure 6D). While average ripple power was identical for light ON and OFF sessions before stimulation (*p* = 0.93, Cliff’s δ=0.06), we observed a reduction in light ON power in the 50 ms following stimulation. Although this fell just short of statistical significance due to a limited number of sessions (N=12; OFF=4, ON=8; p=0.07), effect size analysis revealed a strong trend towards reduction in ripple power (Cliff’s δ=0.69).

Because the relatively high latency precludes intervention during brief events, we focused our analysis of ripple properties on the longest 25% detected ripples by session (Figure 6E, Table S10). With this analysis, we observed substantial alterations of median ripple dynamics induced by optogenetic stimulation. Most notably, light ON sessions displayed significantly lower ripple energy than light OFF (*p* = 0.048, δ=0.75). Additionally, although the reduced sample size and large variation precluded statistical significance for other metrics, effect size analysis revealed strong trends towards neuromorphic-driven optogenetic stimulation inducing a reduced proportion of long ripples (R<80, δ=0.56), higher spectral entropy (δ=-0.56), lower peak frequency (δ=0.56), and higher low frequency contribution (δ=-0.69).

Taken together, these results indicate that closed-loop neuromorphic intervention successfully manipulated prolonged events, resulting in weaker, shorter, and more broadband oscillatory signatures. This spectral shift toward lower frequencies and higher entropy may reflect a desynchronization of the ripple network. Ultimately, despite the latency constraints of the preliminary hardware configuration - which our offline validation confirms can be improved via shorter buffers and direct triggering (Figure 5) - these results provide a successful proof-of-concept that neuromorphic computing can be implemented into closed-loop systems for real-time, state-dependent manipulation of fast brain rhythms *in vivo*.

## 3. Discussion

This study demonstrates a fully integrated neuromorphic closed-loop framework for real-time detection and manipulation of fast hippocampal oscillations, validated for the first time in vivo. By combining compact spiking neural networks, SpiNNaker neuromorphic hardware and the open-source Open Ephys acquisition platform, we implemented a complete sensing-processing-stimulation pipeline capable of detecting hippocampal ripples and triggering feedback stimulation within the temporal window of the ongoing event. The system achieved closed-loop latencies of approximately 50 ms, enabling intra-event intervention in up to 80% of ripples. Neuromorphic-triggered optogenetic inhibition significantly altered ripple dynamics and reduced oscillatory energy, demonstrating effective real-time modulation of hippocampal activity. These results directly address a central gap in neuromorphic neural interfaces: the lack of accessible, in vivo closed-loop systems capable of processing live neuronal data, detecting fast transient biomarkers and driving state-dependent intervention with low latency.

A key objective of this work was to determine whether compact SNNs could detect brief electrophysiological events with sufficient accuracy for closed-loop operation. The trained networks detected ripples across 23 recording sessions with high recall and moderate precision, reaching a median F1-score of 0.56. This performance profile is consistent with the known difficulty of automated ripple and high-frequency oscillation detection, where visually annotated events must be discriminated from sharp transients, muscle artifacts and irregular high-frequency activity with overlapping spectral content (Hagen et al., 2021; Liu et al., 2022; Roux et al., 2017; Zelmann et al., 2012; Zhou et al., 2022). Importantly, the SNNs achieved performance comparable to larger deep learning models while using a substantially smaller architecture, comprising only 41 leaky integrate-and-fire neurons and 530 trainable parameters. This compactness is not simply an architectural convenience - it is essential for deployment on low-power neuromorphic substrates and for future implementations in embedded neural interfaces.

The benchmarking analysis also clarifies the specific value of the neuromorphic approach. Optimized classical filter-based detectors, particularly the Buzsáki-style RMS detector, achieved the highest absolute F1-scores when calibrated to this dataset. Thus, the main contribution of the SNN is not superior offline detection accuracy. Rather, the advantage lies in combining competitive detection performance with compactness, event-driven computation, hardware deployability and stable operation in a real-time closed-loop pipeline. Conventional detectors remain highly effective when thresholds and parameters are carefully tuned, but their performance can degrade substantially under standard out-of-the-box settings. This sensitivity to parameter selection is a practical limitation for chronic or adaptive applications, where recording conditions vary across subjects, sessions and brain states (Fernández-Ruiz et al., 2019; Liu et al., 2022; Navarrete et al., 2016; Roux et al., 2017). In contrast, the SNN maintained competitive performance without post-hoc detector optimization, supporting its suitability for autonomous closed-loop applications.

The feature analysis of detected and missed events further highlights the intrinsic ambiguity of ripple detection. True positives were characterized by stronger ripple-band power, higher peak frequency and denser UP/DN spike representations, whereas false positives and false negatives occupied a highly overlapping low-amplitude feature space. Dimensionality-reduction analyses confirmed that the event classes did not form clearly separable clusters. This suggests that many detection errors occur near the boundary of manual ripple annotation rather than reflecting a simple model failure. The ambiguity is compounded by the fact that the SNN operates on ripple-band activity, whereas manual annotations are based on sharp-wave ripple complexes identified in wider-band signals. Isolated high-frequency ripple-like events may therefore be detected by the SNN but excluded from manual annotations if they lack an obvious sharp-wave component. This issue is not specific to SNNs; it reflects broader challenges in ripple definition, annotation consistency and inter-rater reliability (Buzsáki, 2015; Liu et al., 2022; Spring et al., 2017). Accordingly, the moderate precision observed here should be interpreted in the context of the physiological and annotation ambiguity inherent to continuous hippocampal recordings.

Deployment on neuromorphic hardware was a critical step toward testing whether trained SNNs could operate in a live neuromorphic closed-loop setting. The transition from software simulation to SpiNNaker hardware introduced a modest performance reduction, consistent with the software-to-hardware gap commonly encountered in neuromorphic systems (Pedersen et al., 2024; Schuman et al., 2017). Nevertheless, online SpiNNaker performance remained comparable to the software baseline at the most balanced operating threshold, and the transition from offline continuous processing to online buffered processing had minimal additional impact. These results show that the trained networks could be transferred to neuromorphic hardware without losing the detection capability required for closed-loop use. Equally important, SpiNNaker processing operated in biological real time and reduced power consumption relative to GPU-or CPU-based simulation, supporting the broader premise that event-driven neuromorphic computation is well suited to edge processing in neural interfaces (Covi et al., 2021; Pawlak & Howard, 2025; Yoo & Shoaran, 2021).

The latency analysis identifies the system-level factors that dictate the feasibility of intra-event closed-loop intervention. For fast, transient oscillations, detection accuracy alone is insufficient – feedback must be delivered before the target event terminates. Hardware-in-the-loop validation demonstrated that direct Python triggering, minimal Open Ephys buffers and appropriate firing thresholds substantially reduced round-trip latencies. The fastest configuration achieved median latencies of approximately 50 ms, potentially displaying slight improvements over recent Open Ephys-based implementations (De Sousa et al., 2022; Dutta et al., 2018), and enabled stimulation before ripple termination in up to 80% of events.

Despite this success, our evaluation identified a clear pathway for substantial future optimization. We isolated approximately 9.8 ms of latency overhead directly attributable to USB communication with the host computer. Mitigating this hardware bottleneck, by upgrading to high-speed PCIe communication protocols (Siegle et al., 2017) or transitioning the entire pipeline onto fully integrated System-on a-Chip designs (Sharifshazileh et al., 2020; Yoo & Shoaran, 2021), would meaningfully accelerate intervention times and further maximize the clinical utility of the system.

Crucially, this analysis links neuromorphic detection to functional intervention timing, revealing a highly tuneable latency-performance trade-off: lower thresholds and shorter buffers prioritize sensitivity and intra-event stimulation, whereas slightly higher thresholds and longer buffers improve precision at the cost of additional delay. This flexibility allows the same neuromorphic framework to be adapted to experimental objectives requiring either maximal temporal precision or greater stimulation specificity.

The in vivo experiments provide the decisive validation of the complete framework. In awake, head-fixed mice, hippocampal LFPs were acquired through Open Ephys, encoded into spike trains, processed by SNNs running on SpiNNaker and used to trigger inhibitory optogenetic feedback. Under the optimized low-latency configuration, stimulation occurred within the ripple time window for a substantial fraction of events and significantly altered long ripple dynamics. The reduction in oscillatory energy indicates that neuromorphic-triggered inhibition actively disrupted the ongoing ripple process rather than merely reporting its occurrence. Additional changes in ripple features, including alterations in duration, spectral structure and frequency composition, should be interpreted as coordinated modifications of the underlying oscillatory state. These findings demonstrate that neuromorphic hardware can be used not only for offline neural event detection, but also for causal, state-dependent manipulation of fast brain rhythms in vivo.

Several limitations should be considered. First, although the SNNs achieved competitive performance relative to larger machine-learning detectors, precision remained limited, and false positives would lead to unnecessary stimulation in some closed-loop applications. Future systems could reduce this burden by incorporating reference channels, movement-state gating, multi-channel consensus, adaptive thresholds or online confidence estimates (De Sousa et al., 2022; Dutta et al., 2018). Second, the current networks operated on ripple-band activity from a limited number of datasets and animals. Validation across additional laboratories, recording systems, behavioral states and annotation protocols will be required to establish generalizability. Third, SpiNNaker provides an accessible and flexible neuromorphic platform, but it is not itself an implantable device. Translation to chronic or clinical applications will require miniaturized application-specific hardware integrating sensing, preprocessing, event-driven inference and stimulation control within a low-power system-on-chip. Finally, the present study focused on inhibitory optogenetic modulation of hippocampal ripples as a stringent proof of principle; future work should test whether similar architectures can support bidirectional control objectives, multi-site stimulation and adaptive policy learning.

Despite these limitations, the framework presented here is broadly extensible. Hippocampal ripples provide a demanding benchmark because they are brief, high-frequency events requiring millisecond-scale feedback, but the same principles can be adapted to other electrophysiological biomarkers. In Parkinson’s disease, neuromorphic event detection could support adaptive deep brain stimulation based on beta bursts or other dynamic biomarkers (Oliveira et al., 2023). In epilepsy, SNN-based seizure or high-frequency oscillation detection could provide low-power substrates for responsive stimulation or seizure forecasting (Contreras, Huang, et al., 2024; Ronchini et al., 2026). More generally, event-driven neuromorphic processing offers a promising route toward wearable and implantable neural interfaces that continuously monitor biological signals while minimizing energy consumption.

In summary, this work establishes a practical neuromorphic architecture for low-latency closed-loop control of fast brain dynamics in vivo. By bridging compact SNNs, neuromorphic hardware and widely used open-source neuroscience infrastructure, we demonstrate that live neural signals can be processed in real time to trigger causal feedback within the duration of transient oscillatory events. This provides a scalable foundation for future adaptive neural interfaces built on event-driven, energy-efficient computation.

## 4. Methods

### 4.1. Dataset - Mouse CA1 Hippocampus LFPs

Network training utilized a publicly available *in vivo* electrophysiology dataset (Figshare:117897), originally curated for machine learning-based sharp wave ripple detection (Navas-Olive et al., 2022, 2024). Additional sessions in a similar format were provided to validate the networks on unseen data (Table S1).

Briefly, local field potentials (LFPs) were recorded from the CA1 hippocampus of awake, head-fixed transgenic mice using high-density µ-LED optoelectrodes (32 channels across 4 shanks, 30 kHz sampling rate, 1Hz-5 kHz band) and the Open Ephys system. Refer to the original study for more information (Navas-Olive et al., 2022).

Ground Truth (GT) was established via expert manual annotation. The shank closest to the CA1 region was identified and event onset was defined by the emergence of the first ripple oscillation or sharp-wave onset, and offset by cessation of ripple or return to baseline.

### 4.2. Signal Conditioning and Spike Encoding

Raw LFPs were bandpass filtered with a 4th-order Butterworth filter to isolate the ripple band (100-250 Hz), selected to align with consensus on rodent ripple features (Liu et al., 2022).

To interface the analog LFPs with the SNNs, continuous filtered signals were converted into discrete spike trains using an asynchronous delta modulation scheme, also known as step-forward (SF) encoding (Lichtsteiner et al., 2008), selected for its efficiency, robustness and proven compatibility with HFO identification (Sharifshazileh et al., 2020).

Within this encoding framework, positive (UP) and negative (DN) spikes are generated at time t when the signal amplitude (*x*(*t*)) deviates from a dynamic baseline (*v*_0_) by a defined threshold (*T_base_*):

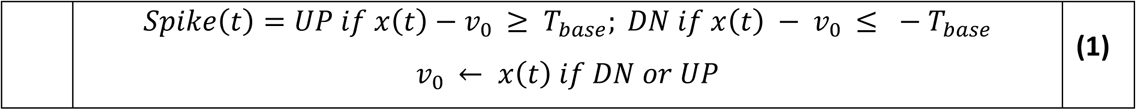

Upon spike generation, *v*_0_ is updated to *x*(*t*). The baseline threshold *T_base_* was calculated dynamically every 10 seconds (50% overlap) based on the background activity of the signal, computed by averaging the peak-to-peak variation of the quietest 25% of 100 ms windows of the baseline segment.

Finally, the 30 kHz spike trains were downsampled to 1 kHz to align with the 1 millisecond timestep required for real-time operation with SpiNNaker (Furber & Bogda, 2020). This was achieved by computing UP/DN net difference over non-overlapping 30 sample (1 ms) windows.

### 4.3. SNN Architecture and Training

Data was segmented into 180 ms windows (120 ms step) to facilitate batch training. Windows were labelled Ripple if they contained the GT ripple onset and extended at least 45 ms beyond it, and assigned a target timing of GT onset + 31 ms (corresponding to the average duration of 4.5 oscillations). Windows without ripples were labelled non-ripple (target=-1, no firing). We removed windows containing fewer than three spikes and applied random undersampling to the non-ripple class to optimize data for training while eliminating the ≈90% class imbalance.

We implemented and trained a simple, fully connected feedforward SNN using snnTorch 0.9.4 (Eshraghian et al., 2023) and PyTorch 2.0.0 (Paszke et al., 2019), optimized for neuromorphic deployment and energy efficiency. All neuronal layers utilized a current-based (CuBa) second-order Leaky Integrate-and-Fire (LIF) model that incorporates synaptic conductance (referred to as “Synaptic” in snnTorch), with dynamics governed by learnable synaptic (α) and membrane (β) decay rates, constrained between 0.01 and 0.99. The network comprised an input layer (UP/DN channels) and two fully connected hidden layers with 24 and 16 LIF neurons, respectively. The second hidden layer is connected to a single-neuron LIF output layer - the “ripple detector” (Figure 1E).

Network weights were initialized via the Xavier normal method (Glorot & Bengio, 2010) with a +0.4 bias applied to positive weights below 0.6 to promote early spiking activity. Synaptic and membrane decay rates were initialized from a normal distribution (*μ* = 0.5, *σ* = 0.2). Training was conducted using Surrogate Gradient Backpropagation Through Time (SG-BPTT) with a fast sigmoid surrogate function (Neftci et al., 2019). Optimization was carried out using the Adam optimizer (Kingma & Ba, 2014), with a base learning rate of 10^−5^, while synaptic weights were updated at a rate five times higher. A StepLR scheduler was employed to decay the learning rate by 5% every 10 epochs.

The training objective minimized a Time-to-First-Spike (TTFS) Mean Square Error (MSE) loss incorporating a 20 ms tolerance window. The loss function was customized to include additional penalizations for FP and FN, which were adjusted throughout training to improve performance.

Models were trained for a maximum of 400 epochs with a batch size of 64. To prevent overfitting, early stopping was applied when no improvement was observed in loss (1% reduction) or accuracy (0.1% increase) over 10 consecutive epochs.

### 4.4. Neuromorphic Closed-Loop System Architecture

#### 4.4.1. System Overview and Data Interface

We developed a fully integrated, real-time closed-loop framework interfacing standard electrophysiology hardware with neuromorphic computing (represented in **Figure 6A**). The end-to-end pipeline consists of four primary stages:

1. **Signal Acquisition**: High-density LFP signals are acquired by an Intan RHD2116 headstage connected via SPI to the Open Ephys acquisition board, which streams data to the host computer running the Open Ephys GUI via USB 3.0 (Siegle et al., 2017).
2. **Software Bridge**: Real-time external access to the data is provided by the Open Ephys ZMQ Interface Plugin. A custom Python application subscribes to this ZeroMQ socket to perform online filtering, spike encoding, and simulation control.
3. **Neuromorphic Processing**: Encoded UP/DN spikes are transmitted via Ethernet to a trained SNN operating in real-time within a SpiNNaker SpiNN-3 board (Painkras et al., 2013).
4. **Stimulation Trigger**: Upon ripple detection, the SNN triggers a low-latency TTL pulse via an Arduino MEGA 2560 to deliver targeted optogenetic stimulation.

#### 4.4.2. Online Preprocessing and Bidirectional Communication

Within the Python software bridge, incoming LFP buffers were continuously filtered using a 4^th^-order Butterworth band-pass filter (100-250 Hz) implemented via SciPy’s *lfilter* function, while preserving the internal filter state (z_i)_ to ensure continuity. Filtered samples were appended to a rolling buffer used to compute the adaptive threshold for spike encoding. Each buffer was subsequently converted into UP/DN spikes and downsampled to match the SpiNNaker timestep, as described above.

The script additionally managed deployment, configuration and real-time interaction with SNN simulations running on SpiNNaker. Bidirectional communication was established via Ethernet using the sPyNNaker “*external_devices”* module (Rhodes et al., 2018), mediated through a *“**LiveSpikesConnection”*** object. Input spikes were streamed to a SpikeInjector population, while output spikes were received in real time from the network’s output neuron and used to trigger immediate closed-loop feedback.

Depending on the experimental configuration, feedback was delivered either as a hardware-triggered TTL via a connected Arduino MEGA 2560 (hardware configuration) or as a software-generated TTL event within the OE GUI using the Network Events plugin. The simulation and data acquisition was configured to run continuously until an external stop command was received.

#### 4.4.3. Neuromorphic Deployment

The trained SNNs were deployed on a neuromorphic SpiNN-3 development board, comprising four SpiNNaker chips (72 ARM968 cores) designed for energy-efficient simulation of large-scale SNNs. The simulation timestep was set to the standard 1 ms, enabling real-time execution (Furber & Bogda, 2020).

Bridging the discrete-time training environment implemented in *snnTorch* and the continuous-time deployment using *PyNN* (0.12.3) on SpiNNaker required systematic adaptations to both neuron model and network architecture. These adjustments ensured that the temporal dynamics of the PyNN model closely matched the trained networks, enabling faithful transfer of learned behaviour to hardware.

Regarding model translation, although both frameworks consist of second-order CuBa LIF neurons, mapping the “*Synaptic*” (snnTorch) model to the “*IF_curr_exp*” (PyNN/SpiNNaker) required several transformations. First, firing threshold and resting potential were scaled for consistency. Second, recursive decay factors (*α*, *β*) were converted into physical time constants ( *τ_syn_*, *τ_m_*), according to *τ* = −1/ ln(*x*).

A further discrepancy arises on the integration of synaptic current in PyNN, which includes a decay scaling factor ((1 − *β*)*R_m_*) absent on snnTorch. To compensate, the membrane resistance *R_m_* was set indirectly by tuning the membrane capacitance *C_m_* to *C_m_* = (*β* − 1)/ln(*β*) to neutralize the decay factor. Finally, excitatory and inhibitory weights were separated into distinct synaptic projections to comply with PyNN’s receptor-type architecture. A fixed 100-millisecond refractory period was imposed on the output neuron to regulate detection frequency, replacing the software-based post-hoc FP suppression.

To preserve the precise temporal structure of the input buffers while conforming to the constraints of the packet-based Address-Event Representation (AER) of SpiNNaker, we implemented a delay-line input architecture. The original 2-channel (UP/DN) input was expanded into a source population whose size matched twice the buffer duration (ms) plus a 10 ms tolerance to accommodate buffer variability and prevent data loss. Spikes occurring at time t within a buffer were routed to specific input nodes assigned a synaptic delay of t+1. Consequently, while all the spikes from a buffer were injected simultaneously into the simulation, they arrived at the first hidden LIF layer sequentially, faithfully preserving the temporal dynamics of the original signal.

#### 4.4.4. Evaluation Configurations

Two configurations were utilized to evaluate the influence of several key parameters, namely firing threshold, buffer and Arduino triggering mode, on system performance:

##### In Silico

Offline streaming of pre-recorded sessions utilizing the Open Ephys File Reader plugin.

##### Hardware-in-the-loop

Emulation of real-time signal acquisition by playing pre-recorded LFPs through a National Instruments DAQ (NI9264). Signals were routed through the I/O interface and acquired by the Open Ephys analog inputs (Figure 5A). LFPs were downsampled to 15 kHz and scaled to ±5 V to match hardware constratints.

During extended hardware-in-the-loop recordings, a minor temporal drift emerged between the DAQ clock and the original recordings. To correct this, we implemented a sliding-window, correlation-based alignment procedure, with the original recording as the temporal reference. Recordings were segmented into 10 s intervals, z-scored, and cross-correlated with the original reference to detect the optimal temporal lag. A linear regression model fitted to these lags yielded the drift slope and the initial offset, which were subsequently applied to the manual annotations to ensure perfectly synchronized latency estimations.

### 4.5. In vivo mice experiments

#### 4.5.1. Animals

All experimental protocols and procedures were approved by the Ethics Committee of the Instituto Cajal and the Spanish National Research Council (CSIC) in strict accordance with Spanish legislation (RD 53/2013 and Law 32/2007) and the European Communities Council Directive (2010/63/EU) on the care and use of laboratory animals. For closed-loop experiments, we used two distinct transgenic mouse lines to target GABAergic interneurons: the PV-Cre line (B6.129P2-Pvalbtm1(cre)Arbr/J (10)) and SST-Cre line (Ssttm2.1(cre)Zjh/J). Animals were housed either individually or in social groups to ensure their well-being. They were maintained under a 12-hour light/dark cycle (lights on at 7:00 a.m.) at a controlled temperature of 21-23°C and 50-65% humidity, with food and water available ad libitum.

#### 4.5.2. Experimental Protocol - Optogenetic Ripple Manipulation

Wideband (1 Hz-5 kHz) local field potential (LFP) signals were pre-amplified (4-10 × gain) and recorded using the Open Ephys Acquisition Board running under the Open Ephys GUI software (Siegle et al., 2017). Electrophysiological recordings and closed-loop optogenetic ripple manipulations were performed using two types of optoelectrodes. First, we used integrated µLED optoelectrodes from Plexon (20 µm site separation, 32 channels and 12 µLEDs spanning 120 µm) to target the entire CA1 pyramidal cell layer. For optogenetic stimulation, selected µLEDs were activated using the Plexon NeuroLight Stimulator (Plexon Inc.) to deliver blue light at low (10-20 nW) and medium (1-10 µW) intensities. Second, we used a Poly3 silicon probe array from Neuronexus (25 μm site resolution, 177 μm2 site area) coupled to an optical fiber. This configuration delivers blue light stimulation in the 100-200 mW range using a diode-pumped solid-state laser (MBL-F-473/300 mW; CNI Laser). The online stimulation protocol consisted on square pulses (100 ms) delivered automatically upon ripple detection.

### 4.6. Performance Evaluation

To evaluate detection performance, we utilized precision, recall, and the F1-score, as these metrics are specifically suited to the inherent class imbalance of ripple events and avoid the ambiguity associated with defining True Negatives (TN) in continuous neural recordings. As the SNNs operated on a single input channel, validation was performed using the same channel employed during acquisition when recordings were obtained with our system. Otherwise, the channel exhibiting the highest ripple-band power was selected (Navas-Olive et al., 2024). Given the current lack of consensus on the optimal evaluation window for ripple detection (Hagen et al., 2021; Liu et al., 2022; Navas-Olive et al., 2022), we adopted a tolerant but consistent criterion for assigning true and false detections.

For offline validation, a True Positive (TP) event was defined when the output neuron spiked within an interval of [-20,100] ms relative to GT ripple onset. Ripples were considered a FN when there were no spikes within this window. FPs were defined as detections outside valid windows and aggregated into 100 ms bins. In addition, each ripple was assigned an extra 100 ms tolerance to emulate a refractory period. The same strategy was applied when benchmarking against alternative detection methods.

For closed-loop/online evaluations, this evaluation window was shifted to [0,120] ms to accommodate system latency, and FP aggregation and extra tolerance were replaced by the refractory period, ensuring consistency. Clinical error rates were expressed as False Positives/Negatives per minute (FPR/FNR). Latencies were calculated relative to ripple onset, and relative latency was calculated considering each ripple’s annotated duration (Dutta et al., 2018).

Event features (Figure 3) were extracted from a [-50,50] ms window centered on the peak ripple power. Ripple power was estimated via the Hilbert transform of the filtered data, squared and smoothed (5 ms kernel), and z-scored across the entire session. Entropy, peak frequency, low frequency contribution and skewness of curvature were calculated using established procedures (Gliske et al., 2020; Navas-Olive et al., 2022). For in vivo ripple manipulation analyses, peri-stimulation power profiles of detected ripples were not z-scored to preserve absolute amplitude dynamics; instead, session-averaged power traces were normalized to the [0,1] range to characterize decay dynamics with stimulation. Other event-related metrics were computed within the [−50,50] ms window around the peak ripple power after z-scoring. Exceptions included event duration, the fraction of long ripples (exceeding 80 ms), and total energy, which were calculated directly from the annotated ripple intervals to preserve their temporal dimension.

Power benchmarks were conducted during inference using the CodeCarbon 3.2.8 Python library (Courty et al., 2026) and the native SpiNNaker energy report.

### 4.7. Statistical Analysis

Statistical analyses were performed in Python 3.12.9 using SciPy 1.15.2 and statsmodels 0.14.6. Non-parametric tests were employed throughout, including the Mann-Whitney U test (independent groups) and Wilcoxon signed-rank test (paired data). The Holm-Bonferroni correction was applied to mitigate effects of multiple comparisons. Correlation analyses were conducted using Pearson’s and Spearman’s correlation coefficients. Statistical significance was defined as *p* < 0.05.

Effect magnitudes were quantified using Cliff’s Delta (δ)(Cliff, 1993) for independent data, interpreted as negligible (δ < 0.147), small (δ < 0.33), medium ( δ < 0.474) and large (δ > 0.474) (Romano et al., 2006). For dependent data, the matched-pairs rank biserial correlation coefficient (thresholds 0.11, 0.28, 0.43) was employed. Dimensionality reduction and visualization was performed via Principal Component Analysis (PCA) and Uniform Manifold Approximation Projection (UMAP) (McInnes et al., 2020).

## Acknowledgments

This work was supported by ‘la Caixa’ Foundation - CaixaResearch Health 2022 (grant HR22-00189). We would like to thank Andrew Rowley and the rest of the SpiNNaker team for providing us with the SpiNN-3 development board and providing technical support over the duration of the project. We would also like to thank Josh Siegle for providing Open Ephys assistance.

