## Supplementary material for "Low-latency neuromorphic closed-loop control of hippocampal ripples *in vivo*": Figure S# / Table S#

### Supplementary Info - Tables

*Table S1 – In vivo mouse recording sessions used during training and validation of the neuromorphic spiking neural network ripple detectors.*

| Animal | Session | Duration (s) | # Ripples |
|---|---|---|---|
| **Training Data** | | | |
| Amigo2 | Amigo2_2019-07-11_11-57-07 | 2398.86 | 1309 |
| Som2 | Som2_2019-07-24_12-01-49 | 1036.25 | 485 |
| Dlx1 | Dlx1_2021-02-12_12-46-54 | 1021.34 | 211 |
| Thy7 | Thy7-2020-11-11_16-05-00 | 744.21 | 1064 |
| **Total** | | **5200.66** | **3069** |
| **Validation Data** | | | |
| (7BBF257) PV13 (1) | 2025-09-22_17-42-27 | 630.40 | 109 |
| (7BBF257) PV13 (1) | 2025-09-22_17-55-26 | 623.99 | 185 |
| (7BBF257) PV13 (1) | 2025-09-23_15-50-26 | 623.26 | 139 |
| (7BBF257) PV13 (1) | 2025-09-23_16-17-52 | 622.29 | 72 |
| (7BBF257) PV13 (1) | 2025-09-24_10-24-40 | 624.79 | 92 |
| (7BBF257) PV13 (1) | 2025-09-24_11-34-51 | 397.89 | 71 |
| (803EF97) PV14 (2) | 2025-09-24_16-29-07 | 637.11 | 40 |
| (7BBF257) PV13 (1) | 2025-09-25_11-21-53 | 622.55 | 202 |
| (803EF97) PV14 (2) | 2025-09-25_16-41-14 | 625.47 | 144 |
| Calbai32 (3) | Calbai32FPGA_251003_144832 | 682.01 | 193 |
| Calbai32 (3) | Calbai32FPGA_251003_150055 | 670.40 | 186 |
| PV01 (4) | PV01ai32FPGA_250611_115923 | 676.00 | 210 |
| PV01 (4) | PV01ai32FPGA_250611_122326 | 700.24 | 169 |
| Calb (5) | Calb_251210_122327 | 657.26 | 91 |
| Calb (5) | Calb_251210_165332 | 666.58 | 91 |
| Calb (5) | Calb_251210_164150 | 654.16 | 118 |
| Calb (5) | Calb_251210_121141 | 650.91 | 69 |
| Calb (5) | Calb_251210_115904 | 694.30 | 56 |
| Calb (5) | Calb_251210_162849 | 702.32 | 113 |
| Calb (5) | Calb_251211_110650 | 651.22 | 68 |
| Calb (5) | Calb_251211_105518 | 650.74 | 131 |
| Calb (5) | Calb_251209_160255 | 349.72 | 66 |
| Calb (5) | Calb_251211_104316 | 665.90 | 104 |
| **Total** | **N = 23, 5 animals** | **14479.51** | **2719** |

*Table S2 – Performance of the SNNs per session (median considering the 12 selected networks). Total represents the median F1, precision and recall over all the sessions, as well as the total sum of TP, FP and FN measurements. Sessions are sorted in a descending order considering the F1-score.*

| Session | F1-score | Precision | Recall | TP | FP | FN |
|---|---|---|---|---|---|---|
| 2025-09-22_17-55-26 | 0.74 | 0.64 | 0.9 | 166 | 95.5 | 19 |
| 2025-09-25_11-21-53 | 0.72 | 0.77 | 0.7 | 142 | 42 | 60 |
| Calbai32FPGA_251003_150055 | 0.71 | 0.59 | 0.9 | 168 | 115.5 | 18 |
| 2025-09-23_15-50-26 | 0.69 | 0.62 | 0.8 | 111.5 | 70 | 27.5 |
| Calbai32FPGA_251003_144832 | 0.69 | 0.57 | 0.85 | 164.5 | 121 | 28.5 |
| Calb_251209_160255 | 0.66 | 0.51 | 0.96 | 63.5 | 60.5 | 2.5 |
| 2025-09-25_16-41-14 | 0.61 | 0.52 | 0.8 | 115.5 | 107 | 28.5 |
| PV01ai32FPGA_250611_115923 | 0.61 | 0.68 | 0.55 | 114.5 | 54 | 95.5 |
| PV01ai32FPGA_250611_122326 | 0.6 | 0.5 | 0.82 | 138 | 138.5 | 31 |
| 2025-09-24_10-24-40 | 0.58 | 0.55 | 0.67 | 62 | 50 | 30 |
| Calb_251210_164150 | 0.57 | 0.41 | 0.92 | 108.5 | 157 | 9.5 |
| Calb_251211_105518 | 0.56 | 0.4 | 0.92 | 121 | 180 | 10 |
| Calb_251210_162849 | 0.55 | 0.39 | 0.94 | 106 | 166.5 | 7 |
| Calb_251210_165332 | 0.55 | 0.39 | 0.94 | 86 | 135.5 | 5 |
| Calb_251210_122327 | 0.52 | 0.37 | 0.89 | 81 | 137 | 10 |
| 2025-09-23_16-17-52 | 0.48 | 0.44 | 0.55 | 39.5 | 48.5 | 32.5 |
| 2025-09-22_17-42-27 | 0.45 | 0.53 | 0.41 | 45 | 42 | 64 |
| Calb_251211_110650 | 0.44 | 0.29 | 0.92 | 62.5 | 148.5 | 5.5 |
| Calb_251211_104316 | 0.44 | 0.28 | 0.95 | 98.5 | 247 | 5.5 |
| Calb_251210_121141 | 0.42 | 0.28 | 0.86 | 59.5 | 154.5 | 9.5 |
| Calb_251210_115904 | 0.39 | 0.24 | 0.94 | 52.5 | 162 | 3.5 |
| 2025-09-24_11-34-51 | 0.31 | 0.59 | 0.23 | 16.5 | 13.5 | 54.5 |
| 2025-09-24_16-29-07 | 0.26 | 0.15 | 0.92 | 37 | 203.5 | 3 |
| **Total** | **0.56** | **0.50** | **0.89** | **2159** | **2649.5** | **560** |

*Table S3 - Summary of median network performance for the 12 networks selected following windowed pre-validation, sorted by descending F1-score. #2 was selected for neuromorphic deployment on the SpiNNaker SpiNN-3 board, due to good performance on preliminary validations.*

| Network | F1-score | Precision | Recall | TP | FP | FN |
|---|---|---|---|---|---|---|
| #1 | 0.61 | 0.61 | 0.70 | 1694 | 1051 | 1025 |
| **#2** | **0.59** | **0.47** | **0.87** | **2071** | **2157** | **648** |
| #3 | 0.58 | 0.47 | 0.89 | 2130 | 2546 | 589 |
| #4 | 0.58 | 0.58 | 0.68 | 1671 | 1318 | 1048 |
| #5 | 0.58 | 0.52 | 0.84 | 2025 | 2016 | 694 |
| #6 | 0.58 | 0.52 | 0.84 | 2014 | 1964 | 705 |
| #7 | 0.55 | 0.5 | 0.89 | 2179 | 2755 | 540 |
| #8 | 0.53 | 0.44 | 0.92 | 2254 | 3135 | 465 |
| #9 | 0.53 | 0.42 | 0.90 | 2240 | 3471 | 479 |
| #10 | 0.51 | 0.43 | 0.92 | 2272 | 3424 | 447 |
| #11 | 0.5 | 0.41 | 0.94 | 2335 | 3956 | 384 |
| #12 | 0.49 | 0.38 | 0.96 | 1694 | 1051 | 1025 |

*Table S4 - Performance summary with different ripple detectors for standard and optimized configurations, reported as median and inter-quartile range (IQR) across sessions. Thresholds reported in the original studies and those found to represent optimal performance are indicated. For Dutta and Buzsaki, this threshold represents the number of standard deviations above the mean, with Buzsaki including high and low thresholds, for detecting and validating candidate events, respectively. As for RippleNet, CNN and LSTM, this represents the probability for ripple detection triggering. For SNN, it represents the firing threshold and was not modified throughout the study, highlighting the relative stability of our approach.*

| | Method | Th | F1 | Precision | Recall |
|---|---|---|---|---|---|
| **Standard** | Buzsaki | 1; 4 | 0.50 (0.37-0.59) | 0.34 (0.25-0.44) | 0.93 (0.88-0.95) |
| | Dutta | 4 | 0.49 (0.33-0.53) | 0.35 (0.22-0.45) | 0.75 (0.67-0.80) |
| | RippleNet | 0.7 | 0.48 (0.43-0.58) | 0.52 (0.40-0.69) | 0.71 (0.58-0.80) |
| | CNN | 0.7 | 0.60 (0.48-0.66) | 0.58 (0.49-0.88) | 0.63 (0.38-0.69) |
| | LSTM | 0.4 | 0.53 (0.39-0.57) | 0.51 (0.36-0.74) | 0.70 (0.48-0.76) |
| | **SNN (this work)** | **1** | **0.56 (0.44-0.63)** | **0.50 (0.38-0.58)** | **0.89 (0.75-0.92)** |
| **Optimized** | Buzsaki | 4; 6 | 0.70 (0.61-0.72) | 0.65 (0.58-0.79) | 0.70 (0.59-0.79) |
| | Dutta | 7 | 0.63 (0.58-0.68) | 0.71 (0.66-0.84) | 0.57 (0.48-0.62) |
| | RippleNet | 0.7 | 0.58 (0.46-0.63) | 0.77 (0.56-0.94) | 0.49 (0.35-0.66) |
| | CNN | 0.7 | 0.60 (0.48-0.66) | 0.58 (0.49-0.88) | 0.63 (0.38-0.69) |
| | LSTM | 0.5 | 0.55 (0.41-0.60) | 0.62 (0.45-0.83) | 0.59 (0.38-0.69) |
| | **SNN (this work)** | **1** | **0.61 (0.53-0.67)** | **0.61 (0.53-0.73)** | **0.70 (0.50-0.79** |

*Table S5 - Electrophysiological characteristics of different network events - True Positives (TP), False Positives (FP) and False Negatives (FN). Data is presented as median (IQR). FN and FP events are compared using Mann-Whitney U test and the Cliff's delta effect size measure.*

| Metric | TP | FN | FP | FN vs FP (p) | FN vs FP (δ) |
|---|---|---|---|---|---|
| **Mean_P (z)** | 2.8 (1.7-4.6) | 1.1 (0.6-1.8) | 1.0 (0.6-1.6) | 0.091 | 0.04 |
| **Peak_P (z)** | 14.4 (9.0-22.5) | 6.5 (4.0-10.4) | 5.79 (3.9-9.0) | 0.004** | 0.07 |
| **H_Freq (Hz)** | 134 (118-150) | 114 (90-136) | 102 (82-122) | 2.14e-15*** | 0.2 |
| **AFR (Hz)** | 320 (250-410) | 180 (140-240) | 230 (170-290) | 8.50e-40*** | -0.33 |
| **Entropy** | 3.55 (3.34-3.75) | 3.65 (3.41-3.86) | 3.59 (3.30-3.82) | 1.81e-05*** | 0.11 |
| **Skew_Curve** | 1.50 (1.23-1.83) | 1.42 (1.14-1.78) | 1.41 (1.11-1.81) | 0.832 | -0.01 |

*Table S6 - Sensitivity Analysis of a selected trained SNN (#2) in ripple detection following SpiNNaker deployment and closed-loop integration. Values are reported as median (IQR), to display the distribution of performance across sessions. The three threshold values considered to display the best results are highlighted in bold.*

| Th | F1 | Precision | Recall | FNR (E/min) | FPR (E/min) |
|---|---|---|---|---|---|
| 0.2 | 0.05 (0.03 - 0.08) | 0.02 (0.02 - 0.04) | 0.76 (0.62 - 0.93) | 1.6 (0.99 - 3.38) | 304.39 (294.52 - 332.2) |
| 0.4 | 0.07 (0.05 - 0.09) | 0.03 (0.02 - 0.05) | 0.91 (0.89 - 0.94) | 0.87 (0.65 - 1.15) | 279.32 (255.24 - 293.95) |
| 0.6 | 0.17 (0.12 - 0.22) | 0.09 (0.06 - 0.13) | 0.91 (0.88 - 0.95) | 1.11 (0.45 - 1.6) | 93.03 (85.57 - 104.13) |
| 0.7 | 0.32 (0.23 - 0.41) | 0.19 (0.13 - 0.26) | 0.91 (0.85 - 0.94) | 0.96 (0.51 - 2.31) | 38.13 (27.55 - 45.02) |
| **0.8** | **0.5 (0.4 - 0.58)** | **0.37 (0.27 - 0.45)** | **0.87 (0.75 - 0.92)** | **1.16 (0.68 - 2.64)** | **16.88 (10.24 - 19.28)** |
| **0.9** | **0.55 (0.48 - 0.65)** | **0.5 (0.39 - 0.6)** | **0.77 (0.56 - 0.83)** | **2.68 (1.29 - 4.7)** | **8.14 (4.72 - 9.56)** |
| **1.0** | **0.56 (0.39 - 0.61)** | **0.62 (0.52 - 0.7)** | **0.56 (0.35 - 0.64)** | **5.01 (3.27 - 7.73)** | **3.46 (2.13 - 4.07)** |
| 1.1 | 0.33 (0.15 - 0.47) | 0.7 (0.44 - 0.72) | 0.23 (0.1 - 0.35) | 7.65 (5.97 - 11.35) | 1.11 (0.97 - 1.56) |
| 1.2 | 0.1 (0.06 - 0.13) | 0.55 (0.45 - 0.65) | 0.06 (0.03 - 0.08) | 9.31 (7.53 - 13.02) | 0.45 (0.29 - 0.71) |
| 1.4 | 0 (0 - 0) | 0 (0 - 0) | 0 (0 - 0) | 10.71 (8.2 - 14.1) | 0 (0 - 0.09) |

*Table S7 - Comparison of the performance of SNN ripple detectors simulated in software (snnTorch), as well as on hardware (SpiNNaker), with both Offline and Online (closed-loop configurations). Results are reported as median (IQR) to display the distribution of performance across sessions. We observed a generalized decrease in F1 following neuromorphic deployment (around 10%). Going from offline to online configuration increased the recall at the cost of reduced precision, improving F1 for higher thresholds ( with a higher precision and lower recall).*

| Method | Th | F1 | Precision | Recall |
|---|---|---|---|---|
| **snnTorch** | 1.0 | 0.59 (0.5 - 0.65) | 0.47 (0.41 - 0.6) | 0.87 (0.67 - 0.92) |
| **Offline** | 0.8 | 0.53 (0.44 - 0.61) | 0.48 (0.36 - 0.56) | 0.77 (0.66 - 0.82) |
| | 0.9 | 0.54 (0.48 - 0.6) | 0.56 (0.44 - 0.62) | 0.7 (0.51 - 0.74) |
| | 1.0 | 0.54 (0.37 - 0.62) | 0.66 (0.57 - 0.75) | 0.51 (0.3 - 0.55) |
| **Online** | 0.8 | 0.5 (0.4 - 0.58) | 0.37 (0.27 - 0.45) | 0.87 (0.75 - 0.92) |
| | 0.9 | 0.55 (0.48 - 0.65) | 0.5 (0.39 - 0.6) | 0.77 (0.56 - 0.83) |
| | 1.0 | 0.56 (0.39 - 0.61) | 0.62 (0.52 - 0.7) | 0.56 (0.35 - 0.64) |

*Table S8 - DAQ Validation Results (Buffer, threshold fixed at 0.8) - Influence of buffer on round-trip latency and performance. Results are expressed as median (IRQ).*

| Buffer (ms) | Latency (ms) | Latency (%) | F1 | Precision | Recall |
|---|---|---|---|---|---|
| 3 | 48.53 (41.37 - 56.52) | 79.45 (62.82 - 97.44) | 0.46 (0.37 - 0.52) | 0.32 (0.24 - 0.4) | 0.84 (0.73 - 0.92) |
| 6 | 51.43 (44.73 - 59.2) | 84.15 (67.8 - 102.11) | 0.5 (0.39 - 0.55) | 0.34 (0.27 - 0.45) | 0.86 (0.73 - 0.9) |
| 10 | 52.28 (45.6 - 60.29) | 86.56 (68.42 - 104.43) | 0.53 (0.4 - 0.59) | 0.37 (0.32 - 0.5) | 0.85 (0.71 - 0.92) |
| 16 | 59 (51.54 - 67.5) | 96.23 (76.4 - 117.77) | 0.5 (0.4 - 0.59) | 0.35 (0.31 - 0.49) | 0.88 (0.76 - 0.92) |
| 20 | 62.47 (55.87 - 69.92) | 102.48 (83.35 - 122.42) | 0.54 (0.44 - 0.63) | 0.43 (0.36 - 0.53) | 0.9 (0.68 - 0.92) |
| 32 | 74.73 (66.83 - 83.03) | 121.42 (99.02 - 144.65) | 0.54 (0.42 - 0.59) | 0.39 (0.33 - 0.51) | 0.87 (0.7 - 0.93) |
| 40 | 81.62 (75.2 - 89.01) | 133.15 (108.53 - 158.08) | 0.57 (0.45 - 0.62) | 0.47 (0.35 - 0.57) | 0.86 (0.73 - 0.92) |

*Table S9 - DAQ Validation Results (selected thresholds) - Influence of buffer and threhsold on round-trip latency and performance in hardware-in-the-loop simulations. Results are expressed as median (IRQ).*

| Th | Buffer (ms) | Latency (ms) | Latency (%) | F1 | Precision | Recall |
|---|---|---|---|---|---|---|
| 0.8 | 3 | 48.53 (41.37 - 56.52) | 79.45 (62.82 - 97.44) | 0.46 (0.37 - 0.52) | 0.32 (0.24 - 0.4) | 0.84 (0.73 - 0.92) |
| 0.8 | 6 | 51.43 (44.73 - 59.2) | 84.15 (67.8 - 102.11) | 0.5 (0.39 - 0.55) | 0.34 (0.27 - 0.45) | 0.86 (0.73 - 0.9) |
| 0.9 | 3 | 52.9 (45.36 - 62.43) | 86 (68.38 - 104.13) | 0.52 (0.39 - 0.57) | 0.41 (0.29 - 0.5) | 0.74 (0.52 - 0.81) |
| 0.9 | 6 | 55.93 (48.57 - 64.43) | 90.28 (72.75 - 108.8) | 0.54 (0.44 - 0.6) | 0.43 (0.33 - 0.56) | 0.76 (0.55 - 0.81) |
| 1 | 3 | 58.63 (51.05 - 67.67) | 91.03 (74.89 - 108.96) | 0.42 (0.28 - 0.51) | 0.4 (0.27 - 0.48) | 0.52 (0.3 - 0.58) |
| 1 | 6 | 61.67 (53.3 - 70.57) | 97.16 (78.63 - 113.42) | 0.47 (0.33 - 0.57) | 0.5 (0.36 - 0.59) | 0.51 (0.31 - 0.62) |

*Table S10 – Event properties of the 25% longest detected ripples across stimulation (Light ON) and no stimulation (Light OFF) sessions (median per session). Metrics include ripple duration (ms), ratio of ripples longer than 80 milliseconds (R>80), peak and mean power (z-scored), peak frequency, entropy, energy and low frequency contribution. Mann-Whitney U probability tests were performed, and Cliff's δ was used to evaluate effect size.*

| Metric | Light OFF (n=4) | Light ON (n=8) | P (MU) | Cliff's δ |
|---|---|---|---|---|
| Duration (ms) | 79.2 (73.20-85.50) | 73.6 (71.70-77.40) | 0.393 | 0.34 (Medium) |
| R>80 | 0.14 (0.09-0.20) | 0.07 (0.04-0.09) | 0.154 | 0.56 (Large) |
| Peak Power (z) | 11.05 (9.63-13.42) | 10.33 (9.92-11.56) | 0.683 | 0.19 (Small) |
| Mean Power (z) | 2.49 (2.43-2.75) | 2.37 (2.13-2.49) | 0.283 | 0.44 (Medium) |
| Peak Frequency (Hz) | 134.5 (127.75-138.25) | 119 (115.75-131.50) | 0.147 | 0.56 (Large) |
| Entropy | 3.51 (3.47-3.59) | 3.64 (3.58-3.72) | 0.154 | -0.56 (Large) |
| Energy | 0.24 (0.23-0.27) | 0.23 (0.20-0.23) | 0.048 | 0.75 (Large) |
| Low Frequency | 0.74 (0.72-0.78) | 0.85 (0.80-0.88) | 0.073 | -0.69 (Large) |

### Supplementary Info - Figures

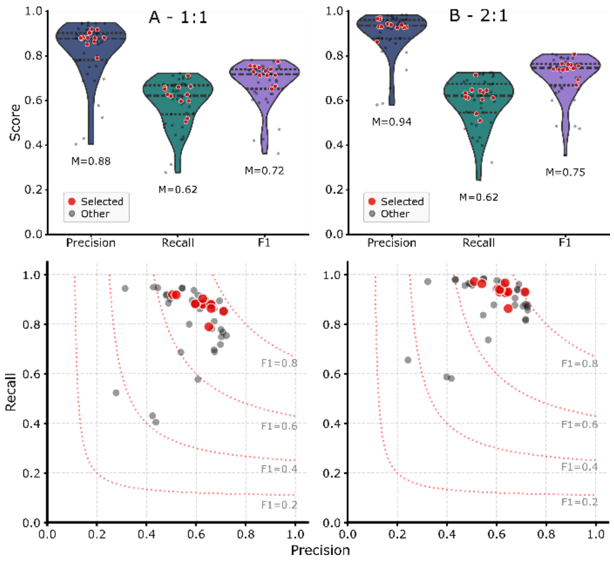


*Figure S1 – Preliminary (class-balanced and windowed) validation of SNNs in ripple detection. Networks selected for further continuous validation and evaluated in the main text are highlighted in red. Windows with 180 ms, annotated similarly to those used for training, were used for preliminary validation. Class balances evaluated were either A. 1:1 (ripple:non-ripple) and B. 2:1. Windows were extracted from the final 20% of each recording session used for training. A True Positive window occurred when the first output spike within the window was between 31 ms before and 45 ms after the Ground Truth annotation, corresponding to the mean and maximum minimal ripple duration calculated previously. Detection outside this range (or missed detections) were False Negatives. Regarding non-ripple windows, those where any spike occurred were classified as False Positives, otherwise they were True Positives.*

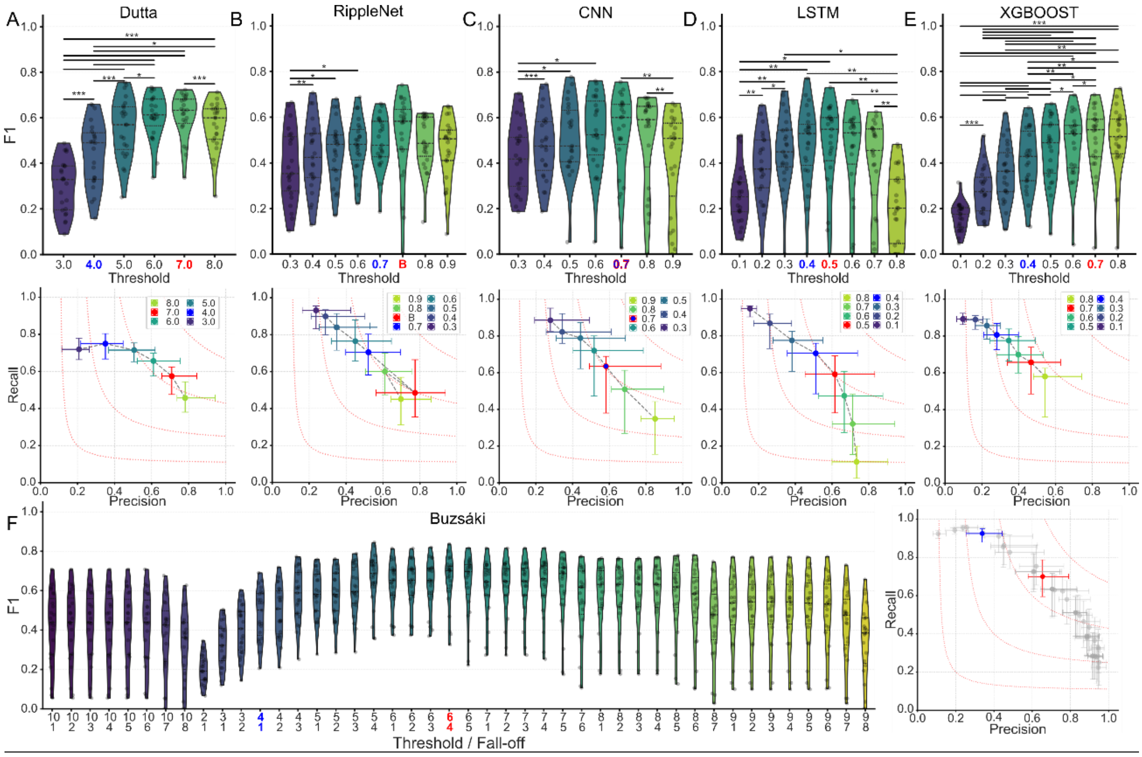


*Figure S2 – Parameter Sweep across different ripple detectors for our dataset (n=23 sessions), to evaluate the performance in standard conditions (here in blue) and find the optimal thresholds for our data (red), obtained by maximizing the F1 for our dataset. Statistical significance is inferred through Wilcoxon signed-rank test, and corrected through the Holm-Bonferroni correction. Annotations correspond to p<0.05 (*), p<0.005 (**), p<0.005 (***). A. Dutta filter. B. RippleNet LSTM. Here, B represents the best model and threshold combination ("unidirectional best_random_seed456", 0.7). C. CNN, with similar standard and optimal parameters. D. LSTM. E. XGBOOST, both from the RipplAI toolbox (Navas-Olive et al., 2024). F. Buzsáki-style RMS-based detector, including threshold and fall-off parameters, that represent the threshold for detection of candidate events and the fall-off for validation regarding event duration.*
